# 30-Color Longitudinal Full-Spectrum Immunophenotyping and Sorting of Human Circulating Immune Cell Subsets Implicated in Systemic Autoimmune Rheumatic Diseases

**DOI:** 10.1101/2025.01.09.631766

**Authors:** Nikhil Jiwrajka, Florin Tuluc, Maria C. Jaimes, Jennifer Murray, Montserrat C. Anguera

## Abstract

This 30-color panel was developed to enable the enumeration and purification of distinct circulating immune cell subsets implicated in the pathogenesis of systemic autoimmune diseases including rheumatoid arthritis (RA), systemic lupus erythematosus (SLE), systemic sclerosis (SSc; scleroderma), Sjögren’s disease (SjD), idiopathic inflammatory myopathy (IIM), and others. While designed for application to peripheral blood mononuclear cells, the inclusion of CD45 coupled with the ability to extract cellular autofluorescence spectral signatures enables the application of this panel to other tissue types. Of the 30 total markers, this panel employs 18 markers to profile T cell subsets consisting of different memory subsets and T helper polarities, ≥ 10 markers to profile B cell subsets including double-negative B cells, and a total of 8 lineage markers to identify immune lineages including monocyte and natural killer cell subsets, conventional dendritic cells, plasmacytoid dendritic cells, and basophils. This panel reproducibly identifies target populations with excellent resolution over several months of data acquisition with minimal batch effects, offering investigators a practical approach to sort immune cell subsets of interest for downstream applications while simultaneously collecting high parameter immunophenotypic information using a limited sample quantity.

## Background

The prevalence and healthcare burden of systemic autoimmunity and systemic autoimmune diseases continues to increase worldwide.^1^ Such conditions, including systemic lupus erythematosus (SLE), rheumatoid arthritis (RA), Sjögren’s Disease (SjD), systemic sclerosis (SSc), and idiopathic inflammatory myopathy (IIM) impart tremendous morbidity upon patients and family members, while also incurring significant healthcare costs that have been estimated to exceed $100 billion annually.^1^ While immunosuppressive therapies can mitigate disease activity, they are often associated with adverse effects, and current therapies do not reliably quell disease activity in all patients for reasons that remain unclear. Additionally, disease activity can also relapse and “flare” in the setting of immune perturbation from myriad causes, including viral infection. While there have been significant advances in our understanding of the immunopathogenesis of systemic autoimmune diseases over the past several decades, the immunological basis of disease development, progression, and especially the risk of disease flare and treatment responsiveness, remains undefined. Animal models have provided valuable mechanistic insights into core immunopathological processes but have been inadequate to pinpoint fundamental disease-specific immunological derangements given that they fail to recapitulate all aspects of human disease. Collectively, these observations highlight the necessity of high-resolution approaches to both comprehensively and deeply profile the immune landscape in well-defined clinical cohorts of humans suffering from these diseases.^2^

The identification of disease-specific immune “signatures” ideally requires the use of high-throughput technologies capable of obtaining data at single-cell resolution. To this end, single-cell transcriptomics have been invaluable in clarifying the complexity of the human immune system. However, these approaches are costly, less readily scalable to large patient cohorts, and furthermore can be prone to undersampling rare, potentially pathologically relevant immune populations, thus requiring complementary approaches to deeply profile the immune system. Conventional flow cytometry has traditionally been an effective complementary modality to study various aspects of human immunology at single-cell resolution. However, the growing appreciation of the complex composition and function of the immune system has led to an expansion of the number of markers required to adequately study this complexity.^3,4^ As a result, comprehensive immune profiling on conventional cytometers has proven challenging, and has often required the simultaneous use of multiple immunophenotyping panels on a single specimen. Such an approach is limited by sample quantity and does not enable a comprehensive assessment of marker co-expression across different cell populations. To this end, the advent of mass cytometry has enabled a remarkable increase in the number of markers that can be measured simultaneously. Despite this major technological advance, mass cytometry also has several logistical limitations, including lower throughput and an inability to sort purified populations for downstream applications because cells are destroyed during data acquisition. Full-spectrum, or “spectral”, flow cytometry builds upon the advantages of conventional flow cytometry and enables an increase in the number of total fluorochromes that can be used in a single panel while simultaneously enabling higher marker resolution, high experimental throughput, and cell sorting. As such, it is well-suited to applications of deep immune profiling of large patient cohorts and can therefore help accelerate new insights into the pathogenesis of systemic autoimmune diseases.

Here, we developed a 30-color spectral cytometry panel for the 5-laser Cytek® Aurora Analyzer and Aurora Cell Sorter to enable a rigorous investigation of the composition and function of the immune landscape in humans with systemic autoimmune disease (**Table 1**). This panel enables the broad identification of innate and adaptive immune cells, and specifically includes markers that permit the identification and purification of immune cell subsets implicated in systemic autoimmune diseases including SLE, SSc, RA, IIM, and SjD (**Table 2**). Though initially developed for application to cryopreserved human peripheral blood mononuclear cells, the inclusion of CD45 also enables comprehensive analysis of tissue-infiltrating immune cells.

**Table 1.**
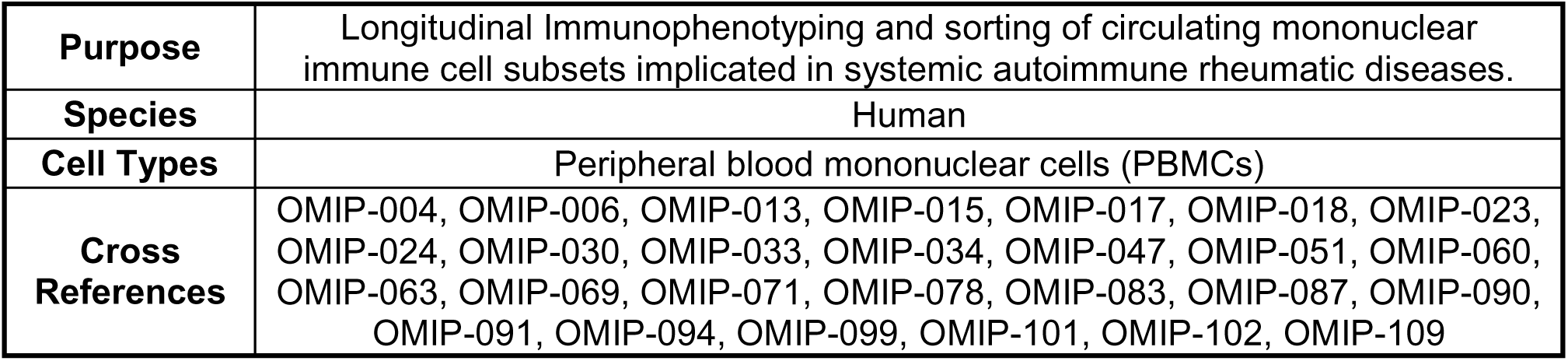
Summary.

|  |  |
| --- | --- |
| **Purpose** | Longitudinal Immunophenotyping and sorting of circulating mononuclear immune cell subsets implicated in systemic autoimmune rheumatic diseases. |
| **Species** | Human |
| **Cell Types** | Peripheral blood mononuclear cells (PBMCs) |
| **Cross**<br>**References** | OMIP-004, OMIP-006, OMIP-013, OMIP-015, OMIP-017, OMIP-018, OMIP-023, OMIP-024, OMIP-030, OMIP-033, OMIP-034, OMIP-047, OMIP-051, OMIP-060, OMIP-063, OMIP-069, OMIP-071, OMIP-078, OMIP-083, OMIP-087, OMIP-090, OMIP-091, OMIP-094, OMIP-099, OMIP-101, OMIP-102, OMIP-109 |

**Table 2.** Reagents used in this panel.

| Specificity | Fluorochrome | Clone | Rationale |
| --- | --- | --- | --- |
| **<u>Viability</u>** |  |  |  |
| **Dead cells** | Live/Dead BLUE | N/A | Viability staining |
| **<u>Leukocytes</u>** |  |  |  |
| **CD45** | PerCP | 2D1 | Leukocyte identification |
| **<u>Monocytes</u>** |  |  |  |
| **CD14** | PE/Fire 810 | S180004C | Monocyte differentiation |
| **CD16** | BV750 | B73.1 | Monocyte and NK cell differentiation; cDC identification |
| **<u>B Cells</u>** |  |  |  |
| **CD19** | BV570 | HIB19 | B cell and antibody-secreting cell (ASC) identification |
| **CD20** | Alexa Fluor 532 | 2H7 | B cell and plasma cell identification |
| **CD21** | BUV661 | B-ly4 | B cell differentiation |
| **CD24** | BUV805 | ML5 | B cell differentiation |
| **IgD** | BUV615 | IA6-2 | B cell differentiation |
| **<u>NK Cells</u>** |  |  |  |
| **CD56** | BV605 | NCAM16.2 | NK cell identification |
| **<u>cDCs</u>** |  |  |  |
| **CD11c** | PE/Cy5 | Bly6 | Conventional dendritic cell identification, B cell differentiation |
| **<u>pDCs</u>** |  |  |  |
| **CD123** | APC/Fire 750 | 6H6 | pDC and basophil identification |
| **<u>T Cells</u>** |  |  |  |
| **CD4** | Alexa Fluor 700 | A161A1 | CD4<sup>+</sup> immune cell subsets (CD4<sup>+</sup> T cells, plasmacytoid dendritic cells (pDC), monocytes, etc.) |
| **CD8** | BUV496 | RPA-T8 | CD8<sup>+</sup> immune cell subsets (CD3<sup>+</sup> T cell subsets, natural killer (NK) subsets) |
| **CD25** | APC | M-A251 | T cell activation and regulatory T cell identification |
| **CD27** | BV785 | O323 | T cell and B cell differentiation and memory state |
| **CD38** | Real Blue 780 | HIT2 | Activation and differentiation of multiple mononuclear immune subsets (T cells, ASCs, and others) |
| **CD45RA** | BUV395 | HI100 | T cell differentiation and memory state |
| **CD95** | BV711 | DX2 | T cell differentiation and memory state, B cell differentiation |
| **CD127** | PE-CF594 | HIL-7R-M21 | CD4<sup>+</sup> and CD8<sup>+</sup> T cell differentiation |
| **CD183**<br>**(CXCR3)** | BV421 | G025H7 | CD4<sup>+</sup> T cell polarity (Th1), T and B cell differentiation |
| **CD185**<br>**(CXCR5)** | BB515 | RF8B2 | T and B cell differentiation |
| **CD194**<br>**(CCR4)** | APC/Fire 810 | L291H4 | CD4<sup>+</sup> T cell polarity (Th2) |
| **CD196**<br>**(CCR6)** | PE/Cy7 | G034E3 | CD4<sup>+</sup> T cell polarity (Th17) and B cell differentiation |
| **CD197**<br>**(CCR7)** | BB700 | 3D12 | T cell differentiation and memory state |
| **CD278**<br>**(ICOS)** | BUV563 | DX29 | T cell activation |
| **CD279**<br>**(PD1)** | BV650 | EH12 | T cell activation, T-follicular and peripheral helper cell identification |
| **CD319**<br>**(SLAMF7)** | PE | 162.1 | Cytotoxic CD4<sup>+</sup> T cell and plasma cell identification |
| **CX3CR1** | BV480 | 2A9-1 | CD8<sup>+</sup> T cell, monocyte, and NK cell differentiation, chemotaxis |
| **HLA-DR** | BUV737 | G46-6 | T cell activation, antigen-presenting cells, pDCs |

Note: Markers are organized in alphanumeric order within each subcategory. Abbreviations:
APC, allophycocyanin; BB, Brilliant Blue; BUV, Brilliant Ultraviolet; BV, Brilliant Violet; PE,
phycoerythrin; PerCP, peridinin-chlorophyll-protein

After excluding dead, CD45^-^ events, monocytes and lymphocytes can be manually identified based on a combination of forward and side-scatter (**Figure 1**). Unbiased analytical approaches can also distinguish monocytes and lymphocytes without the use of forward and side scatter (**Online Figures 10-11**). Monocyte subsets can be broadly distinguished based on expression patterns of CD14 and CD16, which identify classical (CD14^+^CD16^-^), intermediate (CD14^+^CD16^+^), and nonclassical (CD14^-^CD16^+^) monocytes (**Figure 1A**). Of these subsets, classical monocytes are considered to express the highest density of chemokine receptors, thus enabling their chemotaxis to lesional tissues, where they differentiate into macrophages that contribute to inflammatory programs within target organs. Notably, this panel also enables the identification of two more recently described classical monocyte subsets, including SLAMF7^+^ and CD56^+^ classical monocytes. SLAMF7 expression in circulating classical monocytes and tissue-infiltrating macrophages has been associated with immunosuppressive functions in the setting of polymicrobial sepsis^5^ yet hyperinflammatory functions in the context of rheumatoid arthritis.^6^ CD56^+^ classical monocytes also reflect a hyperinflammatory subset of monocytes that are enriched in patients with severe COVID-19 infection^7^ and rheumatoid arthritis.^8^ In contrast to classical monocytes, intermediate macrophages are poised for antigen presentation, and accordingly exhibit greater HLA-DR and toll-like receptor expression.^9,10^ Nonclassical monocytes (NCM) play important roles in endovascular surveillance and possess the ability to adhere to inflamed endothelium via their high expression of CX3CR1, the fractalkine receptor.^10,11^ Notably, NCM have been shown to also migrate towards sites of TLR7-mediated inflammation^11^ and produce proinflammatory cytokines particularly in response to viruses and nucleic acids in a TLR7/TLR8-dependent manner,^12^ thus highlighting their responsiveness to Type 1 interferon-mediated inflammation. These observations are of particular interest in the context of systemic autoimmune diseases including SLE, SjD, SSc, and inflammatory myositis, as many of these diseases exhibit evidence of increased Type 1 interferon signaling in both the circulation and in lesional tissues.^13–15^

**Figure 1.**
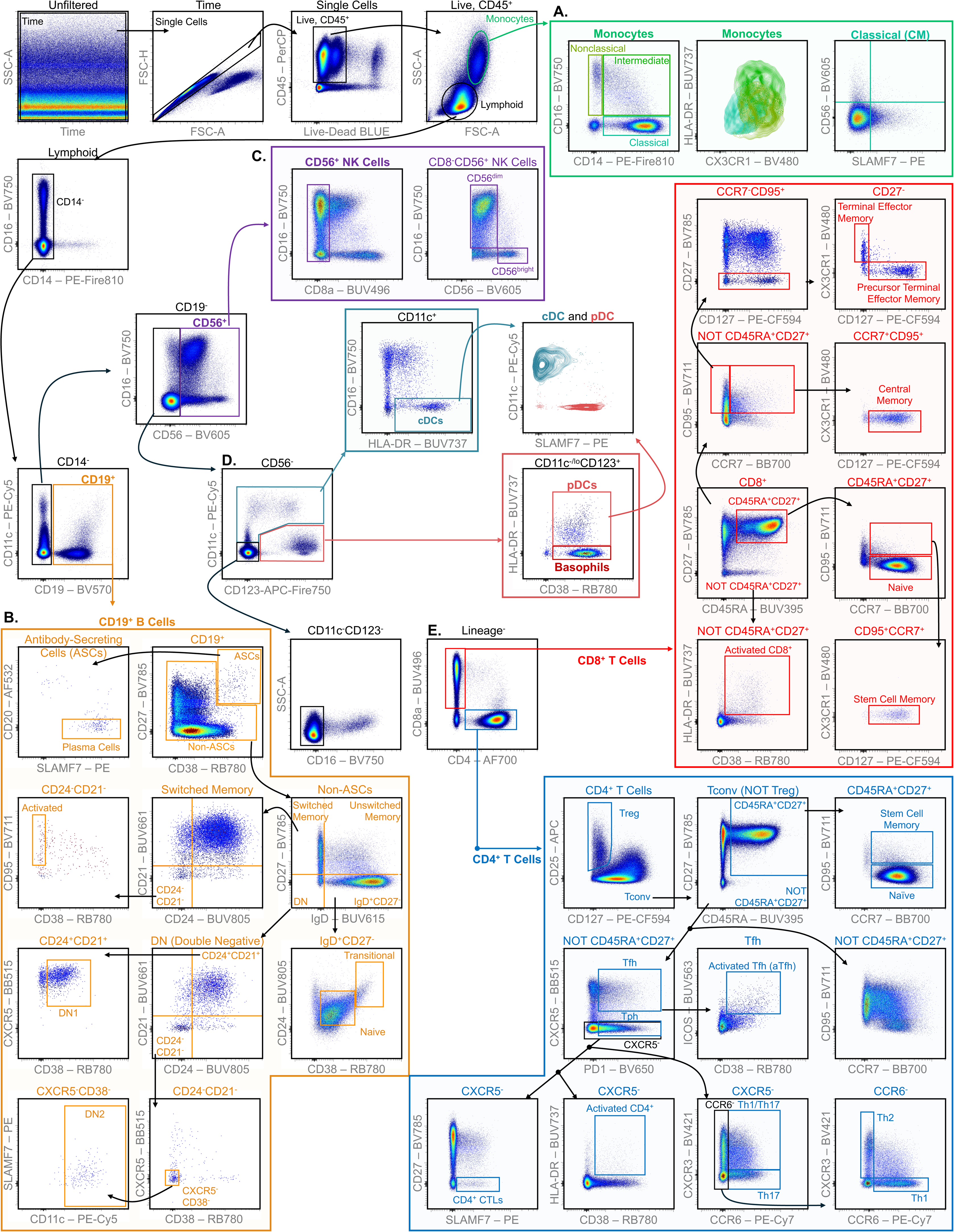
Identification of circulating immune cell populations in cryopreserved PBMCs. Bivariate gating strategy used to identify immune cell subsets of interest. All bivariate plots include concatenated data from n=2 healthy female donors in order to ensure visualization of all cell subsets, as population frequencies vary by donor. All parameters except “Time”, “FSC-A”, “SSC-A”, and “FSC-H”, are arcsinh-scaled using a cofactor of 3000 (PerCP-CD45, BV605-CD56) or 6000 (all other). Cellular subset identification is as follows: **A.** Monocytes (including bivariate plots to depict the relative expression of HLA-DR and CX3CR1, and the proportions of CD56^+^ and SLAMF7^+^ classical monocytes) ; **B.** B cells; **C.** Natural killer cells; **D.** Dendritic cells (including conventional and plasmacytoid dendritic cells, and an additional bivariate plot to depict the relative expression of SLAMF7 amongst dendritic cell subsets); and **E.** T cells (including both CD4^+^ and CD8^+^ T cell subsets).

Amongst lymphoid cells, the B cell lineage can be identified based on the expression of CD19 (**Figure 1B**). Given the abundance of disease-specific, disease-associated, and disease activity-related autoantibodies in systemic autoimmune diseases, and the recent data supporting the efficacy of deep depletion of CD19^+^ B cells^16^, it is clear that the B cell lineage plays an important role in the pathogenesis of multiple systemic autoimmune diseases. Antibody-secreting cells (ASCs) can be identified based on very high expression of CD38 and CD27, and terminally differentiated plasma cells can be further identified based on high SLAMF7 expression and the lack of CD20 expression. Expansions of antibody-secreting subsets have been consistently identified amongst patients with systemic autoimmunity, especially active SLE^17^ and aggressive subtypes of IIM, such as MDA-5 dermatomyositis^18^. Within the remaining B cell compartment, IgD, CD27, CD24, CD38, CD21, CXCR5, CD11c, and CD95 can be used to identify naïve, transitional, founder, unswitched memory (also referred to as circulating marginal-zone or marginal zone-like B cells), class-switched memory, and both DN1 and DN2 B cell subsets, and further distinguish resting from activated B cell subsets.^19^ The activated naïve and DN2 populations are of particular relevance to SLE disease activity and of potential relevance to other diseases including RA and SSc.^17,20–22^ Further characterization of the B cell subset dynamics along the spectrum of disease progression and treatment response may inform our understanding of specific pathogenic B cell subsets in each disease.

The combination of CD56 and CD16, following exclusion of CD14^+^ and CD19^+^ events, enables the identification of human circulating natural killer subsets, including the CD56^bright^CD16^-^ and the CD56^dim^CD16^+^ subsets (**Figure 1C**).^23,24^ These subsets differ with respect to both their abundance in the circulation and their immunologic functions, likely as a result of their unique expression profiles of activating and inhibitory receptors, cytokine and chemokine receptors, and adhesion molecules.^23,25^ Compared to the CD56^bright^CD16^-^ subset, the CD56^dim^CD16^+^ subset is more abundant in the circulation and exhibits greater effector/cytotoxic capabilities, with less proliferative capacity. Deciphering the contributions of these and other NK cell subsets to the immunopathogenesis of systemic autoimmune rheumatic diseases is an area of active investigation. Recent work has implicated activated cytotoxic NK cell subsets in the skin^26^ and lungs^26,27^ of patients with SSc, and in the lungs of patients with myositis-associated lung disease.^28^ Notably, a subset of CD56^+^ NK cells also express CD8. While their contribution to systemic autoimmune diseases is less well studied, circulating CD8^+^ NK cells were found to be inversely proportional to the risk of disease relapse in patients with multiple sclerosis, perhaps via their ability to regulate the proliferation and activation of CD4^+^ T cells.^29^ Future investigation in longitudinal studies may inform their relevance to disease flares in patients with systemic autoimmune diseases.

Of CD56^-^ cells, CD11c, CD16, and CD123 can be used to identify conventional (cDCs; CD11c^+^CD16^-^HLADR^+^) and plasmacytoid dendritic cell (pDCs; CD11c^-^CD123^+^CD38^+^HLADR^+^) subsets (**Figure 1D**). pDCs are thought to play critical roles in the pathogenesis of multiple systemic autoimmune diseases given their propensity to secrete Type 1 interferons and their presence in lesional tissues in multiple diseases.^30,31^ Notably, this panel also reveals that both circulating cDCs and pDCs variably express SLAMF7. In pDCs, SLAMF7 expression can be induced upon *in vitro* stimulation of PBMCs with a combination of SLE-associated autoantibodies and ribonucleoprotein antigens, and its expression is greater amongst interferon alpha-producing pDCs, suggesting that SLAMF7 expression and/or signaling in pDCs may be important in tuning Type 1 interferon responses.^32,33^ Further investigation of pDC abundance and function, and SLAMF7 expression on pDCs in relation to disease activity and responsiveness to immunosuppression in systemic autoimmune diseases characterized by prominent Type 1 interferon-stimulated gene signatures, could help clarify its role in systemic autoimmunity.

Subsequent exclusion of all lineage markers (CD14, CD19, CD56, CD11c, CD123, and CD16) enables the identification of CD4^+^ and CD8^+^ T cells (**Figure 1E**). Notably, because we intended to sort and subsequently activate T cell subsets *in vitro* using anti-CD3/CD28- conjugated beads (Dynabeads^TM^), we did not include CD3 in this panel, as we sought to avoid potential saturation and/or downregulation of CD3, and similarly minimize any functional impact to purified T cells. Despite the absence of CD3 in this panel, ≥99.4% of the remaining CD4^+^ and CD8^+^ cells can be classified as T cells given their co-expression of CD3, as identified via OMIP-069, a distinct high-parameter dataset that shares many of the aforementioned markers (**Online Figure 3**).^34^ In addition to CD4 and CD8, a total of 16 markers, including CD127, CD25, CD45RA, CD27, CCR7, CD95, PD1, CXCR5, CD38, ICOS, HLADR, CX3CR1, SLAMF7, CCR6, CCR4, and CXCR3 can be used to deeply profile CD4^+^ and CD8^+^ T cell subsets implicated in different systemic autoimmune diseases. Notably, this combination of markers identifies a spectrum of memory subsets^35–37^, including T stem cell memory cells (CD45RA^+^CD27^+^CCR7^+^CD95^+^), which represent a highly proliferative memory subset expanded in multiple systemic autoimmune diseases.^38–40^ It also enables the identification of multiple functional subsets that are differentially abundant and/or dysfunctional in active systemic autoimmune disease, including Tregs (CD127^low/-^CD25^high^), T follicular and T peripheral-helper-like cells (CXCR5^+^PD1^+^, CXCR5^-^PD1^+^, respectively), different T helper polarities (based on combinations of CXCR3, CCR4, and CCR6 expression), and effector T cell subsets including activated T cells (ICOS, HLADR, CD38, CD25) and cytotoxic CD4^+^ T cells (CD27^-^SLAMF7^+^).^41–47^

Thus, this panel enables investigators to simultaneously profile multiple immune cell subsets that are highly relevant to the pathogenesis of systemic autoimmune diseases including SLE, RA, SSc, SjD, IIM, and others. The stepwise design and longitudinal application of this panel is described in greater detail in the Online Supplement (**Online Figures 1-10**). Notably, in addition to its ability to broadly profile the innate immune system and deeply profile the adaptive immune system, this panel can also be used without any modifications for the purification of any of the desired immune cell subsets using the Cytek® Aurora CS (**Online Figures 11-12**), thereby enabling downstream studies on purified cell subsets of interest. Thus, we present a unique high-parameter spectral cytometry panel that can be used longitudinally to simultaneously profile and sort immune cells without any adjustments, offering a powerful tool to better understand the immunopathogenesis of systemic autoimmunity, particularly in instances where sample quantity is limited.

## Similarity to Published OMIPs

This panel exhibits similarities to several recently published OMIPs including, but not limited to: OMIP-109 (a recently published 45-color spectral cytometry panel for human PBMC immunophenotyping and sorting using the same instruments described here, with an emphasis on memory T cell subsets)^48^, OMIP-102 (the first 50-color spectral cytometry panel for human PBMC immunophenotyping and cell sorting using the BD FACSDiscover^TM^ S8, with a focus on T cell and DC subsets)^4^, and OMIP-069 (the first 40-color spectral cytometry panel for human PBMC immunophenotyping using the 5-Laser Cytek® Aurora Analyzer)^34^. Our panel’s specific focus on T and B cell subsets implicated in systemic autoimmune diseases, our demonstration of the panel’s robust longitudinal performance for comprehensive immunophenotyping, and our ability to longitudinally analyze and sort cells successfully even using previously recorded reference controls are unique strengths that provide practical utility for investigators studying immune perturbations in systemic autoimmune rheumatic diseases.

## Statement of Ethical Use of Human Samples

De-identified cryopreserved human PBMCs used in this study were obtained from the Human Immunology Core (HIC) Facility at the Perelman School of Medicine at the University of Pennsylvania and from the Benaroya Research Institute (IRB Protocol #10059).

## Author Contributions

Conceptualization: N.J., F.T.

Methodology: N.J.

Data Acquisition: N.J.

Data Curation: N.J.

Data Analysis: N.J.

Validation and Visualization: N.J.

Consultation/Advice: F.T., M.C.J., J.M.

Manuscript Draft: N.J.

Manuscript Review and Editing: N.J., F.T., M.C.J., J.M., and M.C.A

Funding Acquisition: N.J. and M.C.A.

## Acknowledgements

We would like to thank the members of the Human Immunology Core (Max Eldabbas, Emileigh Maddox, Tanishk Sinha, and Jiayi Shu) at the Perelman School of Medicine at the University of Pennsylvania for assistance with sample procurement. We would also like to thank Sylvia Posso and Dr. Jane Buckner from the Benaroya Research Institute for providing several human PBMC samples that were assayed on the Aurora CS. Finally, we would like to thank Dr. Divij Mathew from the University of Pennsylvania for his guidance regarding the Earth Mover’s Distance calculations used to quantify batch effects. This work was supported via grants from the US National Institutes of Health (R21-AR081588 to M.C.A), the Penn Colton Center for Autoimmunity and the Institute for Immunology and Immune Health (to M.C.A. and N.J.), the Rheumatology Research Foundation Scientist Development Award (to N.J.), the Scleroderma Research Foundation Postdoctoral Fellowship (to N.J.), and the H. Ralph Schumacher Rheumatology Research Fund (to N.J.).

## Online Supplement

### INSTRUMENT CONFIGURATION

All flow cytometry analyses described here were performed on a standard 5-laser Cytek^®^ Aurora Analyzer and Aurora CS. The optical layout and instrument setup, is described in detail in the Online Material for OMIP-069.^2^ Information regarding instrument laser and detector modules for the 5 Laser Cytek® Aurora Analyzer and Aurora CS is included in **Online Table 1**. Daily QC using SpectroFlo QC Beads (Lot 2005) was performed on the Aurora Analyzer immediately prior to sample acquisition to help improve consistency across longitudinal sample acquisition. No manual adjustments were made to the manufacturer-recommended assay settings (CytekAssaySetting).

**Online Table S1.**
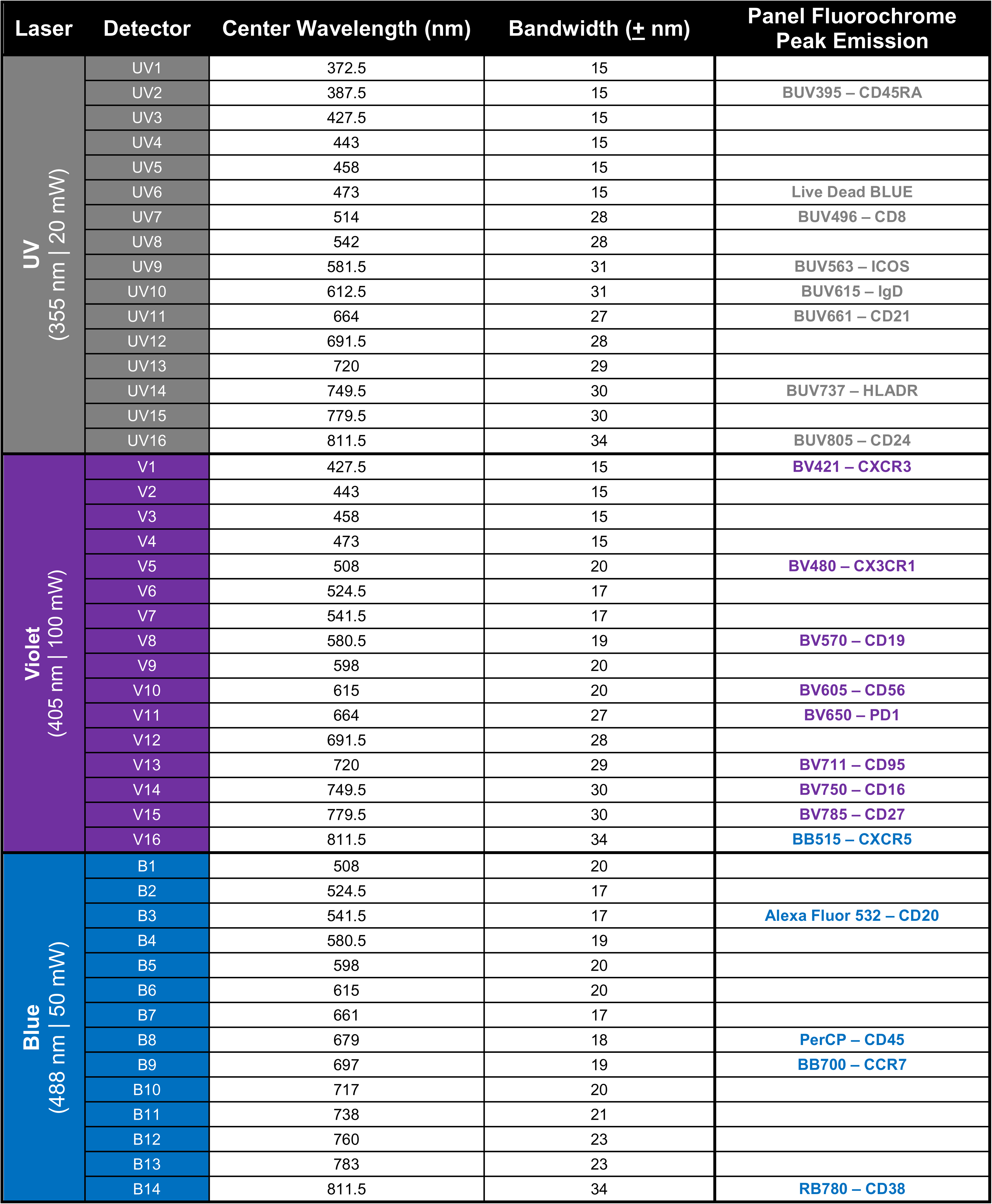
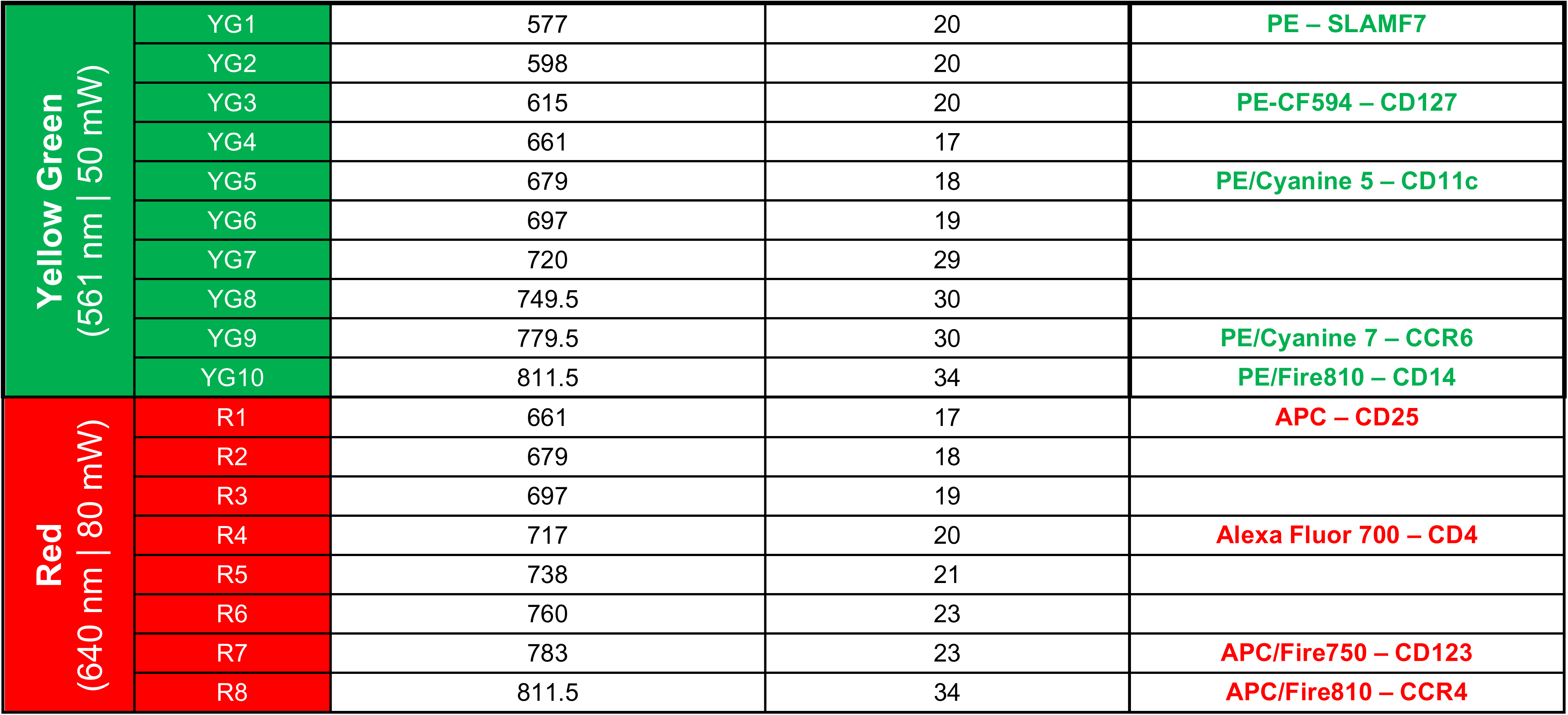
Instrument Configuration for the 5-Laser Aurora Analyzer and Aurora CS. The laser and detector configurations of the Aurora Analyzer and Aurora CS are identical.

| Laser | Detector | Center Wavelength (nm) | Bandwidth (± nm) | Panel Fluorochrome Peak Emission |
| --- | --- | --- | --- | --- |
| UV<br>(355 nm \| 20 mW) | UV1 | 372.5 | 15 |  |
|  | UV2 | 387.5 | 15 | BUV395 – CD45RA |
|  | UV3 | 427.5 | 15 |  |
|  | UV4 | 443 | 15 |  |
|  | UV5 | 458 | 15 |  |
|  | UV6 | 473 | 15 | Live Dead BLUE |
|  | UV7 | 514 | 28 | BUV496 – CD8 |
|  | UV8 | 542 | 28 |  |
|  | UV9 | 581.5 | 31 | BUV563 – ICOS |
|  | UV10 | 612.5 | 31 | BUV615 – IgD |
|  | UV11 | 664 | 27 | BUV661 – CD21 |
|  | UV12 | 691.5 | 28 |  |
|  | UV13 | 720 | 29 |  |
|  | UV14 | 749.5 | 30 | BUV737 – HLADR |
|  | UV15 | 779.5 | 30 |  |
|  | UV16 | 811.5 | 34 | BUV805 – CD24 |
| Violet<br>(405 nm \| 100 mW) | V1 | 427.5 | 15 | BV421 – CXCR3 |
|  | V2 | 443 | 15 |  |
|  | V3 | 458 | 15 |  |
|  | V4 | 473 | 15 |  |
|  | V5 | 508 | 20 | BV480 – CX3CR1 |
|  | V6 | 524.5 | 17 |  |
|  | V7 | 541.5 | 17 |  |
|  | V8 | 580.5 | 19 | BV570 – CD19 |
|  | V9 | 598 | 20 |  |
|  | V10 | 615 | 20 | BV605 – CD56 |
|  | V11 | 664 | 27 | BV650 – PD1 |
|  | V12 | 691.5 | 28 |  |
|  | V13 | 720 | 29 | BV711 – CD95 |
|  | V14 | 749.5 | 30 | BV750 – CD16 |
|  | V15 | 779.5 | 30 | BV785 – CD27 |
|  | V16 | 811.5 | 34 | BB515 – CXCR5 |
| Blue<br>(488 nm \| 50 mW) | B1 | 508 | 20 |  |
|  | B2 | 524.5 | 17 |  |
|  | B3 | 541.5 | 17 | Alexa Fluor 532 – CD20 |
|  | B4 | 580.5 | 19 |  |
|  | B5 | 598 | 20 |  |
|  | B6 | 615 | 20 |  |
|  | B7 | 661 | 17 |  |
|  | B8 | 679 | 18 | PerCP – CD45 |
|  | B9 | 697 | 19 | BB700 – CCR7 |
|  | B10 | 717 | 20 |  |
|  | B11 | 738 | 21 |  |
|  | B12 | 760 | 23 |  |
|  | B13 | 783 | 23 |  |
|  | B14 | 811.5 | 34 | RB780 – CD38 |
| Yellow Green<br>(561 nm \| 50 mW) | YG1 | 577 | 20 | PE – SLAMF7 |
|  | YG2 | 598 | 20 |  |
|  | YG3 | 615 | 20 | PE-CF594 – CD127 |
|  | YG4 | 661 | 17 |  |
|  | YG5 | 679 | 18 | PE/Cyanine 5 – CD11c |
|  | YG6 | 697 | 19 |  |
|  | YG7 | 720 | 29 |  |
|  | YG8 | 749.5 | 30 |  |
|  | YG9 | 779.5 | 30 | PE/Cyanine 7 – CCR6 |
|  | YG10 | 811.5 | 34 | PE/Fire810 – CD14 |
| Red<br>(640 nm \| 80 mW) | R1 | 661 | 17 | APC – CD25 |
|  | R2 | 679 | 18 |  |
|  | R3 | 697 | 19 |  |
|  | R4 | 717 | 20 | Alexa Fluor 700 – CD4 |
|  | R5 | 738 | 21 |  |
|  | R6 | 760 | 23 |  |
|  | R7 | 783 | 23 | APC/Fire750 – CD123 |
|  | R8 | 811.5 | 34 | APC/Fire810 – CCR4 |

### FLUOROCHROME SELECTION AND PANEL DESIGN STRATEGY

This panel was initially designed with the intent of identifying and sorting T cell subsets via CD3 negative selection. Thus, we sought to achieve high resolution of T cell subset populations as well as adequate resolution (and exclusion) of non-T cells. We initially selected antibody-fluorochrome pairs targeting the following 18 markers (CD4, CD8, CD127, CD25, CD45RA, CD27, CCR7, CD95, CXCR5, PD1, CD38, ICOS, CCR6, CXCR3, CCR4, SLAMF7, CX3CR1, HLA-DR) to identify T cell subsets of interest. A gating strategy was established using these markers (**Figure 1E**). This gating strategy was used to inform fluorochrome assignment, with strict attention to minimizing spectral overlap between the XY pairs of a bivariate plot (**Online Figure 1A**). Fluorochrome assignment was also based on established best practices for flow cytometry panel design.^3^ More specifically, we sought to the pair bright fluorochromes with lowly expressed antigens and dim fluorochromes with highly expressed antigens. To gauge relative fluorochrome brightness, we used the publicly available “Cytek^®^ Aurora Fluorochrome Selection Guidelines 5 Laser 16UV-16V-14B-10YG-8R”. Antigen expression profiles were often determined experimentally. However, on several occasions, we analyzed the publicly available FCS files from OMIP-69 for information regarding antigen density and staining pattern (with knowledge of the relative brightness of the fluorochrome from the aforementioned guide).^2^ We preferentially attempted to assign dim fluorochromes (BUV395, BUV496) to highly expressed antigens (CD45RA, CD8), and bright fluorochromes (PE, PE-CF594, BV421, PE-Cy7) to lowly-expressed antigens (SLAMF7, CD127, CXCR3, CCR6, respectively). Selected antibody-fluorochrome conjugates were titrated to determine the optimal concentration (**Online Figure 2A**). This process was pursued iteratively with different antibody-fluorochrome conjugates and clones.

The calculation of the spillover-spreading matrix (SSM) using the statistics from SpectroFlo® software or the OMIQ cytometry analysis platform can be time-consuming. To expedite the evaluation of spillover spreading during earlier stages of panel design, spillover-spreading was assessed based on manual inspection of NxN plots of multicolor samples, with specific attention to the bivariate plots of fluorochrome pairs with high similarity in the spectral emission profile (≥ 25%). Antibody-fluorochrome pairings and staining concentrations, particularly for these culprit fluorochrome pairs (e.g., PE-Cy7-CCR6 and RB780-CD38) were adjusted to (a) maintain adequate population resolution; (b) achieve saturation, particularly for secondary and tertiary antigens; and (c) minimize spillover spreading (**Online Figure 2B**). The following antibody-fluorochrome pairs were tested but ultimately replaced for the specified reasons.

1. **BB515-CCR7** (Clone: 2-L1-A) was eventually replaced with **BB700-CCR7** (Clone: 3D12): Relatively poor separation of the positive and negative populations compared to BB700-CCR7. Furthermore, CCR7 is not highly expressed, and we therefore opted to assign a more highly and widely expressed antigen to BB515 given that it exhibits minimal spectral overlap with other fluorochromes.
2. **BV605-SLAMF7 (**Clone: 235614) was eventually replaced with **PE-SLAMF7 (**Clone: 162.1): Relatively poor separation of the positive and negative populations compared to PE-SLAMF7.
3. **PE/Fire 700-CCR4** (Clone: L291H4) was eventually replaced with **APC-Fire810-CCR4** (Clone: L291H4): Inappropriate initial pairing based on antigen density (highly expressed) and fluorochrome brightness (too bright), resulting in significant spillover spreading into BB700-CCR7. Continued use required reagent dilution well below the saturating concentration. PE-Fire700 was ultimately excluded from the panel to avoid spillover-spreading into BB700.
4. **APC/Fire810-CD38** (Clone: HIT2) was eventually replaced with **Real Blue 780-CD38** (Clone: HIT2): Adequate separation/staining, but CD38 was ultimately moved to Real Blue 780 in order to accommodate CCR4-APC/Fire 810 (limited conjugate availability for CCR4). Notably, Real Blue 780 was not commercially available during the early stages of the panel design process in early 2022.

The purification of T cells via CD3 negative selection requires the inclusion of lineage markers (Lin) to ensure that nearly all Lin^-^CD4^+^ or Lin^-^CD8^+^ cells are indeed CD3^+^ T cells. Thus, we concurrently incorporated 6 additional lineage markers (CD14, CD16, CD11c, CD56, CD19, and CD123), while accounting for the previously noted considerations with antibody-fluorochrome pairing (**Online Figure 1B**). We took care to ensure that each lineage marker exhibited minimal spectral overlap with another lineage marker, and furthermore, that each lineage marker did not receive substantial spillover spreading from the existing T cell subset markers. When spectral overlap between lineage markers was inevitable, we took particular care to appropriately pair fluorochrome brightness with antigen density (example: CD19-BV570 exhibits a discrete positive and negative population of maximal intensity < 10^5^, thereby conferring minimal spillover spreading into CD56-BV605; the same applies to CD4-AF700 into CD123-APC/Fire750). We took note of which of the existing markers were specific to T cells, and were more tolerant of fluorochrome similarity between lineage-defining antibody-fluorochrome conjugates and these

T-cell-specific markers, with the rationale that any resulting spillover-spreading error (from a lineage marker into a T-cell subset marker) would not impart drastic analytical consequences, as this error within T cells would be obviated by the exclusion of non-T-cells when analyzing T cell subsets. As an example, though PE-Fire810 (CD14) exhibits high similarity (69%) with PE-Cy7 (CCR6) and is therefore likely to contribute spillover-spreading into CCR6-PE-Cy7, the resulting spillover-spreading does not ultimately impact the resolution of CCR6 within lymphocytes, the cell types that predominantly express CCR6, as lymphocytes are CD14-negative. Similarly, CD14 exhibits a “primary” pattern of antigen expression, and is therefore its resolution is unlikely to be influenced by mild spillover-spreading of CCR6-PE-Cy7. The following antibody-fluorochrome pairs were tested but ultimately replaced for the specified reasons.

5. **PE-CD56** (Clone: MY31, BD Pharmingen): Relatively poor resolution of CD56^dim^ and CD56^bright^ populations compared to clone **BV605-CD56** (Clone: NCAM16.2, BD Horizon), especially due to spillover-spreading from PE-Fire810-CD14, which partially obscured the resolution of the CD56^dim^ population.
6. **BV510-CD123** (Clone: 9F5, BD Horizon): Relatively poor resolution of CD123^dim^ events compared to **APC-Fire750-CD123** (Clone: 6H6). Resolution was also partially reduced due to spillover spreading from the autofluorescence (AF) parameter upon AF extraction. AF extraction was helpful in improving the resolution of fluorochromes emitting in the V5- V10 and YG1 regions. **BV510** was accordingly not used in the panel.
7. **eFluor450-CD123** (Clone: 6H6, ThermoFisher): Use of this antibody-fluorochrome conjugate resulted in an unusual population of CD123^+^ high side-scatter cells (presumed monocytes) (**Online Figure 3**). This pattern of staining could not be corrected using Fc block products from either BioLegend or BD Biosciences, and furthermore could not be corrected using the True-Stain Monocyte Blocker^TM^ product from BioLegend. We did not use the Fc block product provided by ThermoFisher, and it is possible that this would have resolved the issue. Notably, a similar finding was observed using eFluor450-CD19 (Clone: HIB19), thus suggesting non-specific binding of this fluorochrome to monocytes. This binding pattern was not observed using BV570-CD19 (Clone: HIB19). The same non-specific binding pattern was initially observed with the use of APC/Fire750-CD19 (Clone: HIB19), but subsequently resolved upon addition of the True-Stain Monocyte Blocker^TM^ product from BioLegend.

Finally, given the well-established role of B cells in the pathogenesis of systemic autoimmune rheumatic diseases, we elected to also include specific markers to more deeply profile B cell subsets. While several important markers had already been included (CD27, CD38, CXCR3, CXCR5, CD11c, SLAMF7), we also opted to include several additional markers (CD20, CD21, CD24, IgD). We employed similar principles as discussed above, with special attention to the minimization of spectral overlap between B cell markers (**Online Figure 1C**). We took note of which of the B cell markers of interest were exclusively expressed on B cells, and therefore allowed these B-cell-specific antibody-fluorochrome conjugates to receive spectral overlap from other antigens that are not expressed on B cells, Nevertheless, some degree of spectral overlap (BUV737, BUV805) was inevitable given the limited availability of commercially available spectrally distinct fluorochromes.

The above process was performed iteratively to identify the optimal antibody-fluorochrome pairing and antibody-fluorochrome concentrations, ultimately leading to the final 30-color panel (**Online Figure 4**). The spillover-spreading matrix (SSM) reveals higher spillover-spreading coefficients within the expected fluorochrome pairs. Nevertheless, these fluorochrome pairs maintain adequate population resolution.

**Figure S1.**
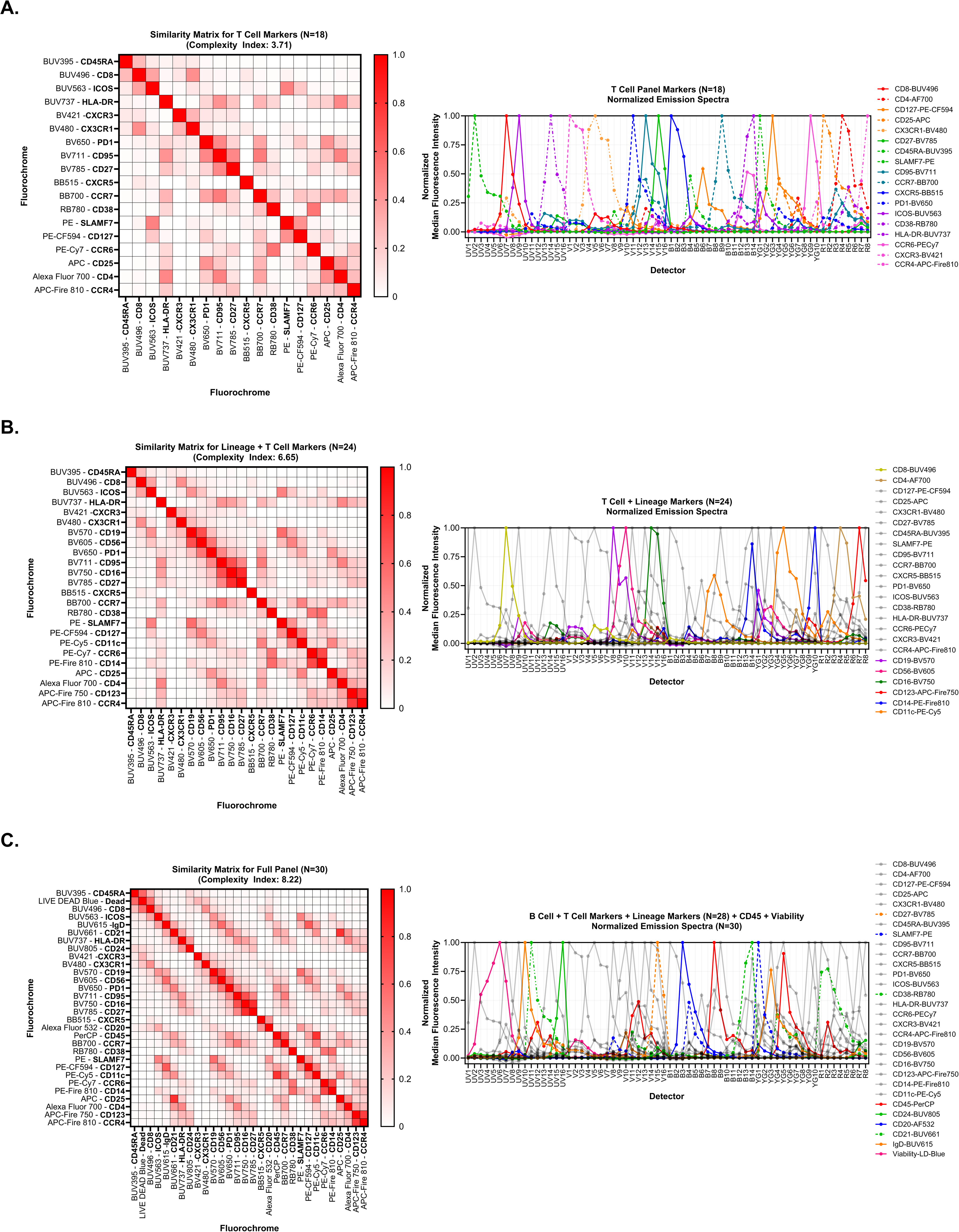
Stepwise approach to fluorochrome and antibody-fluorochrome conjugate selection for immunophenotyping human peripheral blood mononuclear cells. This panel was created in three broad steps, starting with an 18-marker panel to profile CD4^+^ and CD8^+^ T cell subsets. Additional lineage-defining markers, followed by B cell subset markers, were sequentially added to produce the final 30-marker panel. **A**. Similarity indices as calculated in Cytek® Full Spectrum Viewer and normalized emission spectra of n=18 total markers used to identify circulating CD4^+^ and CD8^+^ T cell subsets. XY pairs of a bivariate plot (solid or dashed line) within the gating strategy depicted in the main text share the same color. If a third marker (dot-dashed line) is used in conjunction with either of the XY pairs, it also shares the same color as the XY pairs. **B.** Similarity index and normalized emission spectra of n=6 lineage markers (colored) in addition to the existing n=18 T cell-subset markers (gray). **C**. Similarity index and normalized emission spectra of n=4 additional markers to enable deeper immunophenotyping of B cell subsets (colored, solid or dashed), the viability dye (colored, solid), and the CD45 antibody-fluorochrome conjugate (colored, solid). XY pairs of a bivariate plot (solid or dashed line) within the gating strategy depicted in the main text share the same color. Complexity indices reported in the Cytek^®^ Full Spectrum Viewer are shown. Normalized emission spectra for all n=30 markers were calculated using single-color (cell) controls from the same experiment.

**Figure S2.**
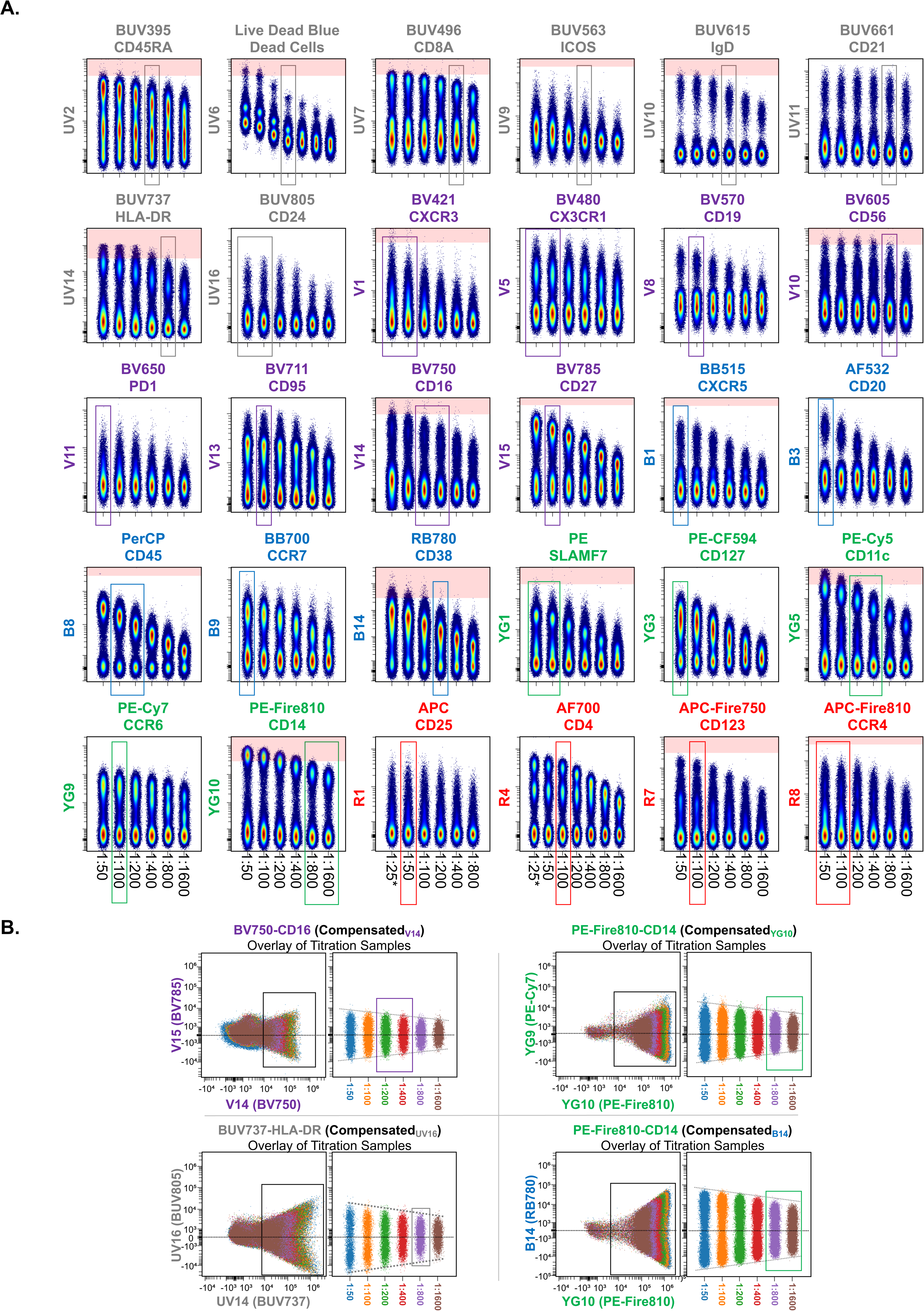
Identification of the optimal concentrations for the selected antibody-fluorochrome conjugates. **A.** PBMCs were stained with a given antibody-fluorochrome conjugate at 6-7 different concentrations (1:50, 1:100, 1:200, 1:400, 1:800, 1:1600; for APC-CD25 and Alexa Fluor 700-CD4, 1:25* was also included). Pseudocolor contour plots of live cellular events (pre-gated on live, single cells identified via FSC-A vs. FSC-H) are depicted for each antibody-fluorochrome conjugate. Each contour plot depicts the detector corresponding to the peak fluorochrome emission (Y-axis) and the staining pattern for each concentration (X-axis; concentration decreases from left-to-right). The “optimal” concentration is depicted using a vertical rectangle (rectangles encompassing two concentrations indicate that the optimal chosen concentration lies in between the depicted concentrations). Fluorescence intensities exceeding 300,000 are highlighted in red, as intensities beyond this threshold were generally associated with more significant spillover-spreading. Conjugate concentrations resulting in staining intensities of this magnitude were avoided due to increased spillover spreading, as depicted in Panel B. **B.** Depiction of the concentration dependence of spillover spreading (introduced from the fluorochrome on the X-axis into the fluorochrome on the Y-axis) for four fluorochromes (X-axis) with high spectral overlap with other fluorochromes included in this panel. The first of the two plots for a given antibody-fluorochrome conjugate is a dot-plot-overlay of cells stained with increasing concentrations of reagent. The second of two plots for a given antibody-fluorochrome conjugate more clearly depicts the concentration dependence of the spillover spreading introduced into the recipient fluorochrome (Y-axis).

**Figure S3.**
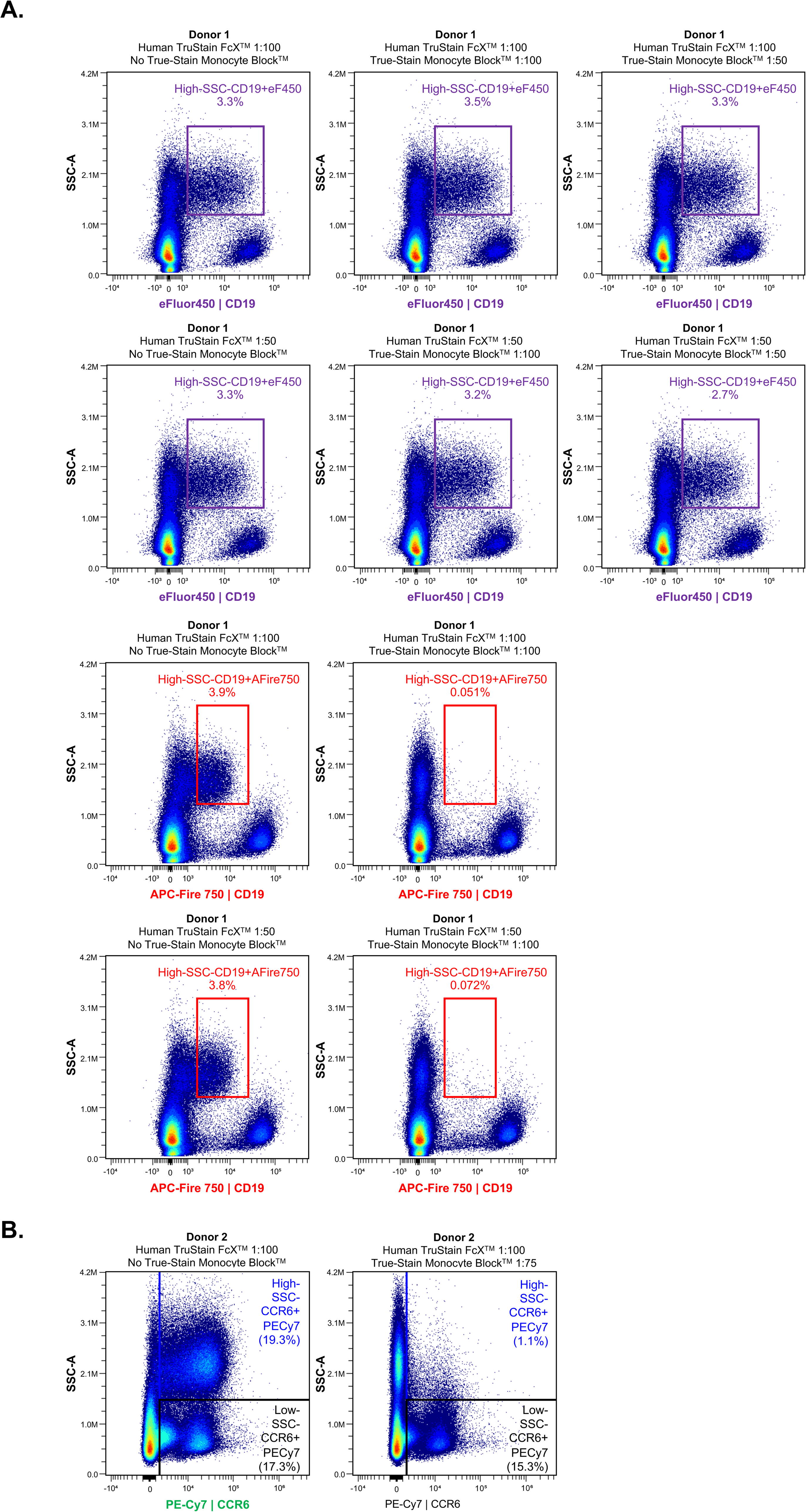
TrueStain Monocyte Blocker^TM^ is required for accurate immunophenotyping. **A.** PBMCs from the same healthy human donor were stained with either anti-human CD19- eFluor^TM^ 450 (Clone: HIB19; purple) or anti-human CD19 APC/Fire^TM^ 750 (Clone: HIB19; red). Cells were stained in the presence of Human TruStain FcX^TM^ at either 1:100 or 1:50, as it is generally accepted that blocking Fc receptors is necessary to mitigate non-specific binding of antibody-fluorochrome conjugates. During the surface stain, cells were then stained in the absence or presence of True-Stain Monocyte Blocker^TM^, an additional reagent that is specifically designed to block the non-specific binding of cyanine-containing dyes, including PE/Cyanine 7, PE/DazzleTM 594, PE/Cyanine 5, Fire dyes, and even BB700.^1^ An aberrant population of CD19^+^ events with high side scatter is apparent in samples stained with both eFluor^TM^ 450 and APC/Fire^TM^ 750. This population is lost with the addition of True-Stain Monocyte Blocker^TM^ to samples stained with APC/Fire^TM^ 750, but persists in samples stained with eFluor^TM^ 450. **B**. PBMCs from a different healthy human donor were stained with anti-human CCR6 PE-Cy7 (Clone: G034E3). In the absence of True-Stain Monocyte Blocker^TM^, there is an abundant high side scatter population (blue gate outline, blue text) that is lost with the addition of True-Stain Monocyte Blocker^TM^.

**Figure S4.**
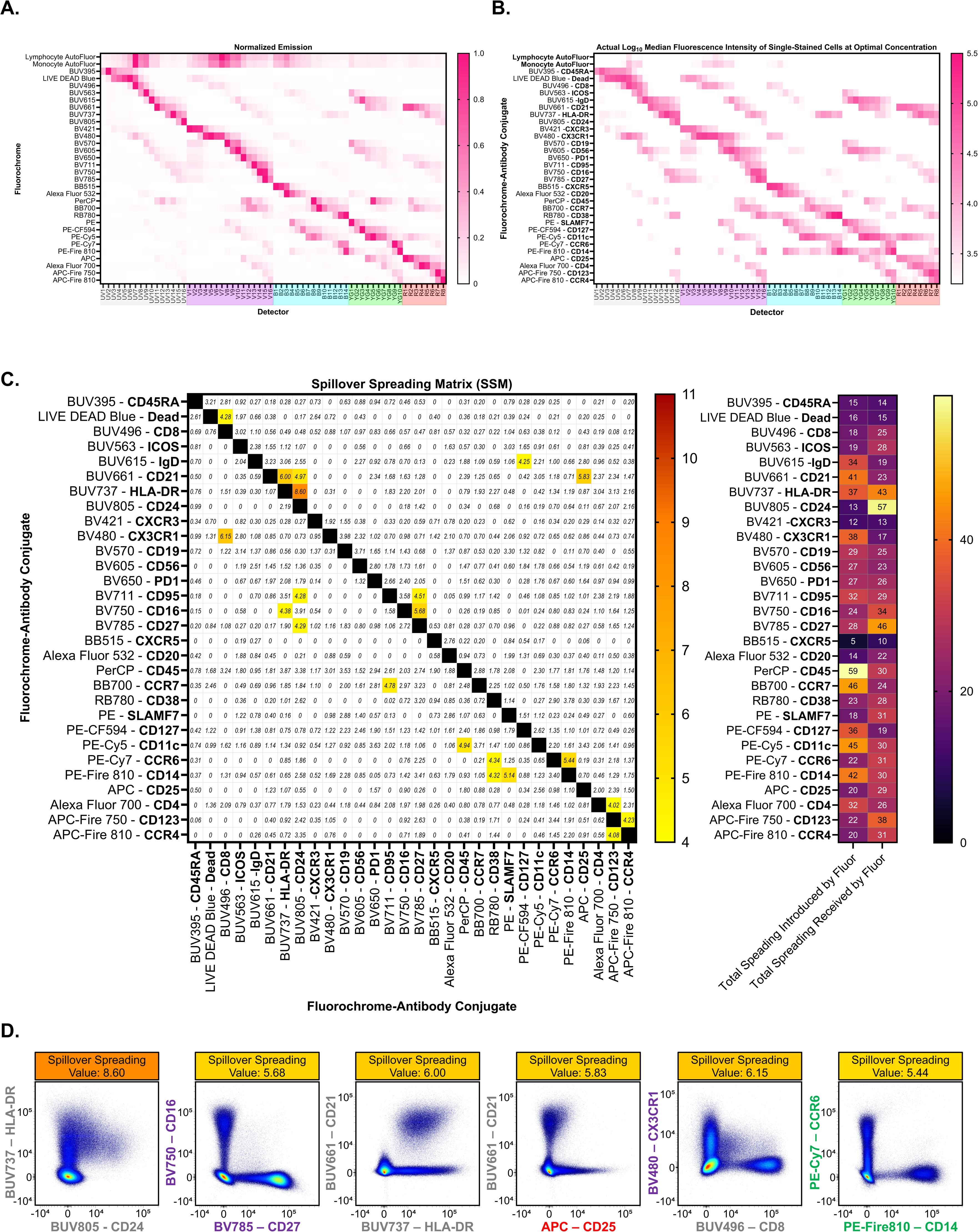
Assessment of spectral overlap and spillover spreading of the finalized panel. **A.** Heat map of the row-normalized emission for each antibody-fluorochrome conjugate based on its median fluorescence intensity on live, single-stained cells at the optimal antibody-fluorochrome conjugate concentrations. **B**. Heat map of the actual median fluorescence intensity (log_10_-scale; MFI) for each antibody-fluorochrome conjugate on live, single-stained cells at the optimal antibody-fluorochrome conjugate concentrations. Deviation of the actual MFI heatmap (panel B) from the idealized MFI heatmap (panel A), particularly along columns (detectors) of highly similar fluorochromes, can help identify fluorochrome pairs where spillover spreading may impact marker resolution (particularly when there is significant overlap in their emission spectra, and that overlap occurs in the peak emission wavelength of one or both fluorochromes). **C.** Spillover Spreading Matrix (SSM) for the fully stained panel. The spillover spreading value refers to a measure of the spillover spreading error introduced by the conjugate in a given row into the conjugate in a given column. **D.** Bivariate plots of select conjugate pairs exhibiting high spillover spreading values (parent population consists of “singlets” identified via FSC-A vs. FSC-H and without exclusion of “Dead” or CD45^-^ events).

### ASSESSMENT OF MARKER RESOLUTION AND PERFORMANCE OF THE FULL PANEL

We subsequently verified that the fully-stained panel, in comparison to single-stained cells, maintains adequate resolution of each marker (**Online Figure 5**). Quantification of the proportion of positive events for each marker confirms adequate performance of the fully stained panel despite some decrease in marker resolution (**Online Figure 5B**). Notably, the largest discrepancies in the proportion of events (≥ 4%) between the single- and multicolor specimens were mostly observed among fluorochromes where single-color stains (stained in the absence of TrueStain Monocyte Blocker^TM^) resulted in a greater proportion of high side-scatter positive events, indicating that these discrepancies are largely due to the non-specific binding of these fluorochromes (BB700, PE-CF594, PE-Cy5, PE-Cy7, PE-Fire810, APC-Fire750, and APC-Fire810) to monocytes (**Online Figure 3B**).^1^

We also sought to confirm that the fully stained samples can accurately identify circulating immune cell subsets of interest. We therefore compared the proportions of various immune cell subsets identified using our panel with those identified via reanalysis of the .fcs files from OMIP-069 using the gating strategy depicted in **Figure 1** (**Online Figure 6**).^2^ Notably, we did not include CD3 in this panel, as our intent was also to devise a panel that would enable cell sorting of T cell subsets via negative selection. We confirmed the appropriateness of our approach using the data from OMIP-069, demonstrating that ≥99.4% of the single or double-positive CD4+ or CD8+ cells resulting from this gating strategy are indeed CD3+ (**Online Figure 6A**).

**Figure S5.**
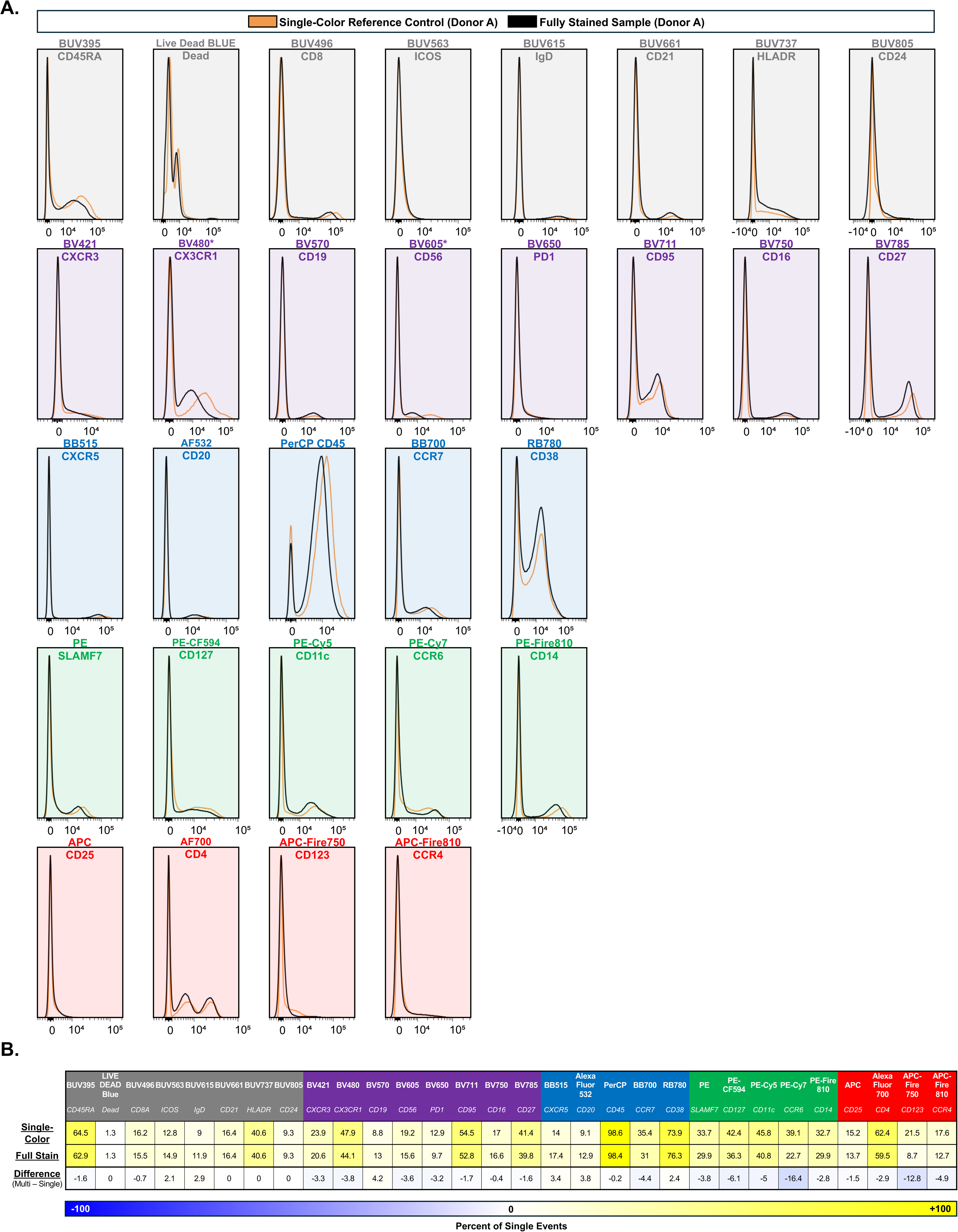
Comparison of marker resolution between single-stained versus fully-stained cells. **A.** Histogram overlay of single-stained (orange) and fully-stained (black) cells from the same donor, using the optimal antibody-fluorochrome concentrations (parent gate comprised of singlets identified via FSC-A vs. FSC-H). Single-stained controls were stained in the presence of TruStain FcX^TM^ but not in the presence of True-Stain Monocyte Blocker^TM^. **B.** Quantitation of positive events, expressed as a percentage of singlet (FSC-A vs. FSC-H), cellular (based on FSC-A vs. SSC-A) events, for the single-stained and fully-stained samples, as well as the absolute difference in the resulting percentages.

**Figure S6.**
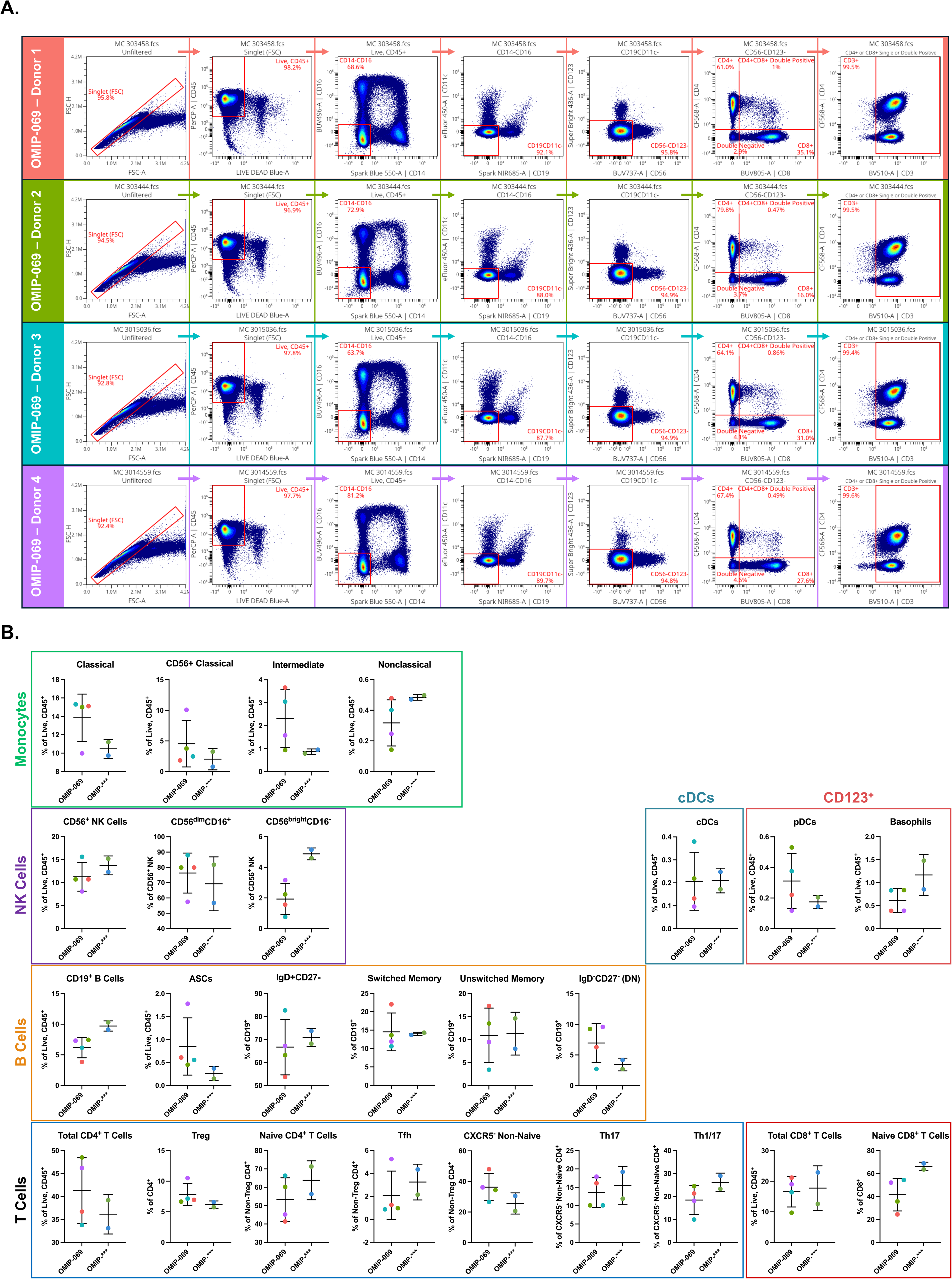
Confirmation of gating strategy to enable sorting of CD4^+^ and CD8^+^ T cells via negative selection. Because we intended to sort and subsequently activate T cell subsets *in vitro* using anti-CD3/CD28-conjugated beads, we did not include CD3 in this panel. To confirm that the majority of the CD4^+^ and CD8^+^ cells identified via the gating strategy depicted in the main text were also CD3^+^ (i.e., T cells), the 4 fully stained samples from OMIP-069 were reanalyzed using the six major lineage markers (CD14, CD16, CD19, CD11c, CD123, CD56) used in our panel. **A.** Analysis of each of the four multicolor samples (MC 303458, MC 303444, MC 3015036, MC 3014559) featured in OMIP-069. Percentages are relative to the parent gate. **B.** Population frequencies of immune cell subsets identified in both the 2 fully stained samples depicted in the main text (Figure 1) and the 4 fully stained samples from OMIP-069. Data are depicted as mean percentages ± standard deviation. Individual multicolor samples from OMIP-069 (n=4) are colored based on panel A. Individual multicolor samples from our panel (n=2) are based on their identity (“Bridge 1” = light green; “Bridge 2” = light blue).

### REFERENCE CONTROL SELECTION

Having confirmed the appropriate performance of the full panel, we also sought to compare the accuracy of spectral unmixing using single-stained cells and single-stained beads. Because reference controls must possess a maximal fluorescence intensity that that is either equivalent to or brighter than the fully stained sample, we first compared the maximal fluorescence intensities of single-stained cells versus single-stained beads (**Online Figure 7A**). This approach identified two antibody-fluorochrome conjugates for which single-stained cells resulted in a higher maximal fluorescence intensity than single-stained beads at the identified “optimal” dilution:

1. CD38-RB780 (likely due to the presence of plasma cells, which are CD38^bright^)
2. CD56-BV605 (likely due to the presence of CD16^-^CD56^bright^ natural killer cells)

These observations indicated that cell-based single-stained controls may be necessary for at least these two antibody-fluorochrome conjugates. We then systematically compared the normalized emission spectra of single-stained cells and single-stained beads **(Online Figure 7B**). While the general shape of the normalized emission spectra appeared to be similar upon cursory visual inspection (with the exception of Alexa Fluor 700-CD4), comparison of the emission spectra at higher resolution revealed that these spectra were not identical, therefore demonstrating that even very small magnitude differences (ΔNormalized MFI_cells-beads_ ≥ or ≤ 0.020) in the normalized emission spectra can result in inaccurate spectral unmixing. The threshold of this difference appears to be proportional to the maximal staining intensity of the antibody-fluorochrome conjugate. These data suggest that conjugates exhibiting high fluorescence intensities (in our experience on the Aurora Analyzer, specifically intensities approaching or exceeding 10^5^) may exhibit erroneous unmixing with another fluorochrome (“Fluor B”) based on the type of reference control, particularly if the absolute value of the difference between reference control emission spectra in the peak detector of “Fluor B” is greater than or equal to 0.020. The threshold for the magnitude of the difference may be higher for antibody-fluorochrome conjugates exhibiting a lower maximal fluorescence intensity (for example: CD127-PE-CF594 versus CD56-BV605).

Given that very subtle differences between the normalized emission spectra of single-stained cells and single-stained beads could contribute to errors in spectral unmixing, we opted to use single-stained cells for all reference controls except CCR6-PE-Cy7. We used single-stained beads for CCR6-PE-Cy7 because (1) the emission spectra of beads and cells do not differ substantially and (2) this conjugate is prone to non-specific monocyte binding that requires use True-Stain Monocyte Blocker^TM^. As a result, when using single-stained cells in the absence of True-Stain Monocyte Blocker^TM^, the “positive” population selected for unmixing should exclusively contain lymphocytes, as the inclusion of any monocytes results in an aberrant emission spectrum that results in spectral unmixing errors (**Online Figure 7B**). We elected to use beads as the reference control for this conjugate to avoid the necessity of using the True-Stain Monocyte Blocker^TM^ reagent for reference control preparation and to ensure appropriate unmixing of CCR6^bright^ events.

**Figure S7.**
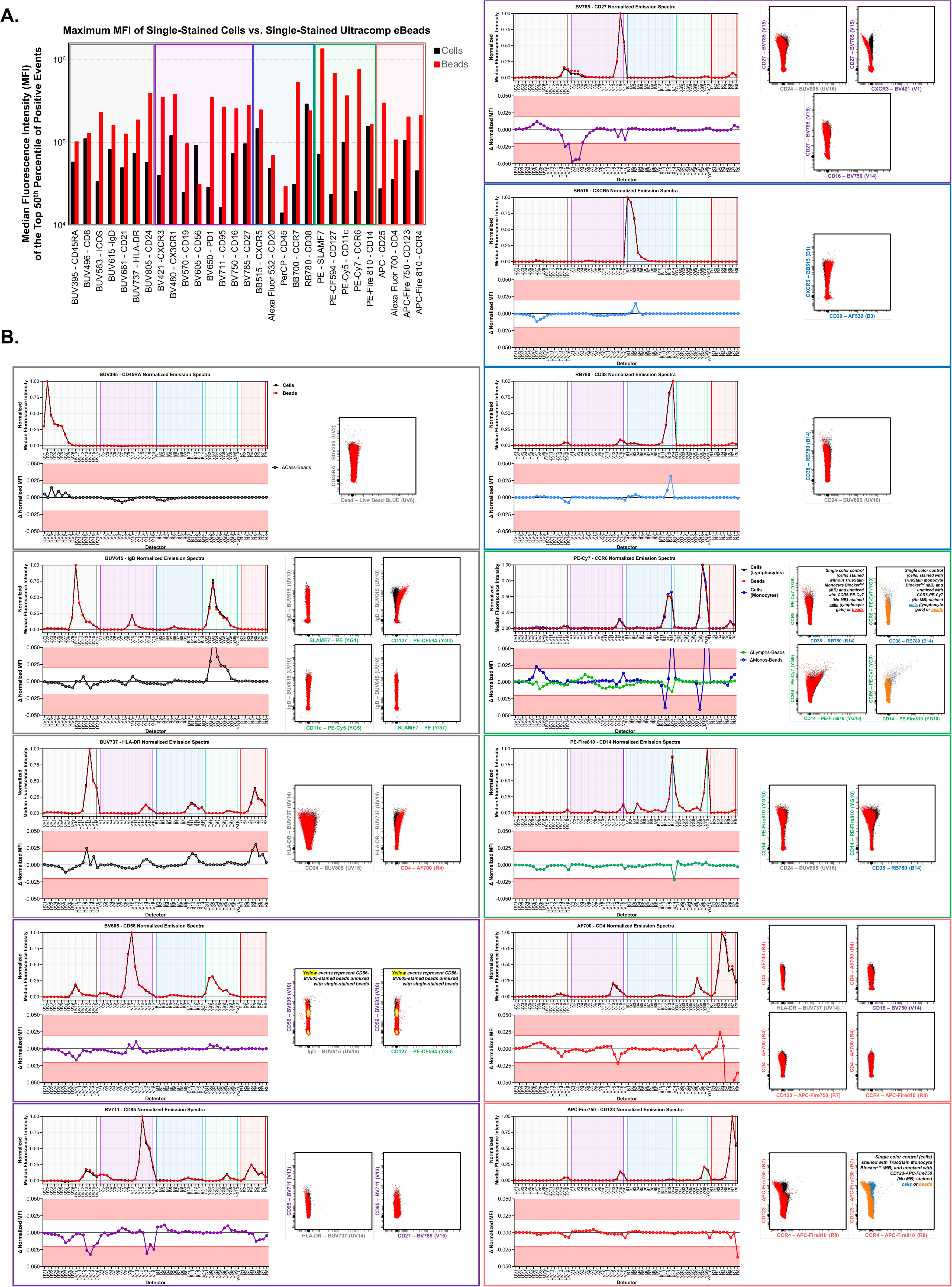
Comparison of reference controls comprised of single-stained cells versus single-stained beads. **A.** Median fluorescence intensities (MFI) of the top 50^th^ percentile of the positive population of single-stained cells and single-stained Ultracomp eBeads^TM^ Compensation Beads for each antibody-fluorochrome conjugate, stained and acquired during a single experiment. These comparisons demonstrate the necessity of using cell reference controls for CD56 and CD38 in particular, given that the fluorescence intensities for these conjugates are higher in single-stained cells than in single-stained beads. **B.** Overlay of the normalized emission spectra of single-stained cells (black, solid) and beads (red, dashed) stained and acquired during the same experiment, as well as the corresponding subtraction plots (ΔNormalized MFI_cells-beads_). Select bivariate plots of single-stained cells unmixed with cells (black) or beads (red) are provided to depict the effects of discrepancies in the emission spectra between single-stained cells and single-stained beads on spectral unmixing. Unmixing errors are appreciable even in the presence of small differences (≥ 0.02 or ≤ −0.02) within the emission spectra between single-stained cells and beads. Based on these observations, single-stained cells were used for all reference controls except for CCR6-PE-Cy7. For this particular conjugate, the brightest population using a single-color control that does not contain True-Stain Monocyte Blocker^TM^ may be incorrectly gated on the aberrant high side-scatter population shown in **Online Figure 3B** (blue, “monocytes”). The normalized emission of the “monocyte” population (blue) is distinct from that of that lymphocyte population (black), and this results in erroneous unmixing. To circumvent this issue, single-stained beads were used for CCR6-PE-Cy7.

### SPECTRAL UNMIXING STRATEGY

The gating process to select the “positive” and “negative” populations within single-stained cells for spectral unmixing impacts the accuracy of spectral unmixing. Because accurate unmixing (as well as compensation) requires that the autofluorescence signature of the “negative” gate be identical to that of the “positive” gate”, we took specific note of the FSC-A and SSC-A coordinate range corresponding to the brightest “positive” events for each single-stained cell control (**Online Figure 8A**; the FSC-A coordinates can be identified using a similar approach). This is because FSC-A and especially SSC-A appear to be positively correlated with autofluorescence within human PBMCs. Such an approach therefore helps to ensure that the autofluorescence signatures of the “positive” and “negative” populations of a given reference control are more homogenous, ultimately resulting in more accurate spectral unmixing. When a “negative” population exhibiting the same range of FSC-A and SSC-A as the “positive” population is not readily identifiable (i.e., CD14-PE-Fire810, CD45-PerCP), a universal negative comprised of unstained cells subsetted to these coordinates can be used instead (**Online Figure 7B**). This approach was used to improve the accuracy of spectral unmixing in the setting of concomitant autofluorescence extraction (**Online Figure 7C**).

**Figure S8.**
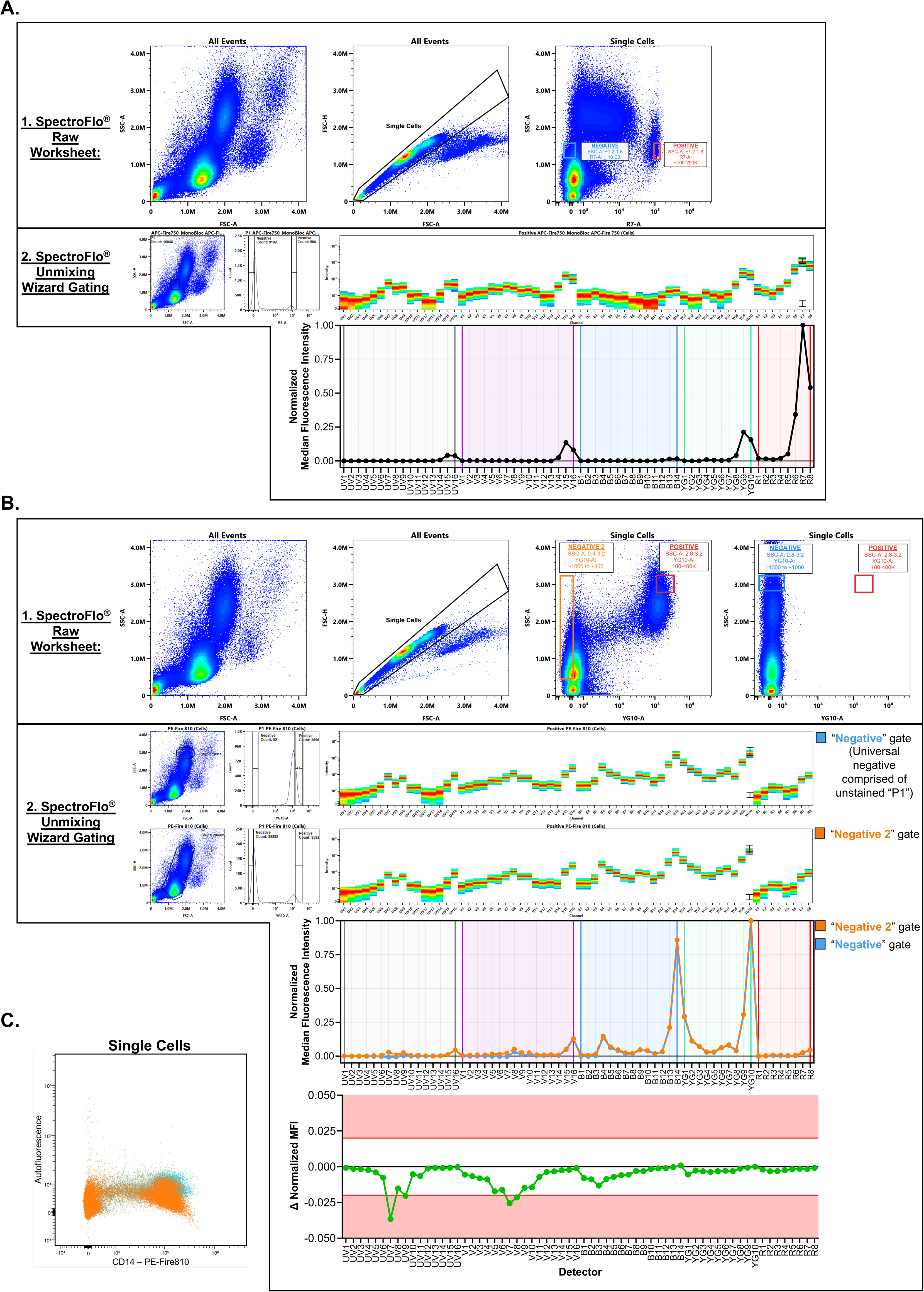
Spectral unmixing strategy. **A.** Example of the approach used to perform spectral unmixing using single-stained cells (stained with APC/Fire750-CD123 containing TrueStain^TM^ Monocyte Blocker). The brightest events among single cells were identified within the SpectroFlo^®^ raw worksheet. To ensure that the autofluorescence signatures of the positive and negative gates ultimately used for spectral unmixing would be similar, we assumed that PBMCs sharing similar FSC-A and SSC-A profiles would tend to exhibit more homogenous autofluorescence signatures, and accordingly identified the specific FSC-A/SSC-A coordinates of the brightest positive and negative populations. These coordinates were then used to identify the positive and negative populations within the SpectroFlo^®^ unmixing wizard. **B.** Another example of the approach used to perform spectral unmixing using single-stained cells (stained with PE/Fire810-CD14). Again, the brightest events among single-stained cells were identified within the SpectroFlo^®^ raw worksheet. In this example, however, there are not sufficient “Negative” events bearing same SSC-A coordinate as the “Positive” events. In such cases we performed spectral unmixing using two different approaches. In one approach, we used a negative population encompassing a large range of SSC-A values (“Negative 2”, orange). In another approach, we used a (universal) negative population comprised of a subset of the unstained sample consisting only of the population exhibiting high SSC-A (“Negative”, light blue). The normalized emission spectra from these approaches are largely similar, but differ subtly along the cellular autofluorescence spectrum. These differences are sufficient to produce erroneous unmixing when cellular autofluorescence is extracted.

### LONGITUDINAL IMMUNE PROFILING AND CONSIDERATIONS FOR REUSE OF PREVIOUSLY RECORDED REFERENCE CONTROLS SPECTRAL UNMIXING

We sought to use this panel to immunophenotype a cohort of human donors (n=74). However, we also intended to sort T cell subsets from each of these donors. Because we only had one aliquot of PBMCs per donor, we devised an approach that would enable us to obtain 30-parameter analytical data while also sorting the T cell subsets of interest (**Online Figure 9A**). Accordingly, these samples were organized into a total of 9 experimental batches for simultaneous application to the 5 Laser Aurora Analyzer and Aurora CS. Each experimental batch consisted of 8-10 samples and 2 bridge samples. Due to the relative ease of use (sample loading, computational speed of unmixing, SpectroFlo® software) of the Aurora Analyzer, we decided to conduct the immunophenotypic analysis using data collected on the 5 Laser Cytek^®^ Aurora Analyzer (as opposed to exclusively using the Aurora CS). Nevertheless, all data was also collected on the Aurora CS.

Because it would be relatively less resource-intensive to reuse previously recorded reference controls (as opposed to preparing new single-color controls for every single experimental batch), we performed monthly experiments (“Control Load”) in which a full set of single-stained cells and beads were reloaded on both the 5-Laser Cytek® Aurora Analyzer and Aurora CS (n=122 samples in total per experiment, including single-stained cells and beads (n=60), and unstained cells (n=1), for each instrument (n=2)). Nevertheless, we quickly noted subtle yet nontrivial variations in the emission spectra even within these 1-month intervals despite our best attempts to account for the many sources of batch-to-batch variation (**Online Figure 9B**). We specifically took great care to ensure identical staining practices, reagent lots, time between staining and sample recording, and instrument setup and quality-control. Ultimately, however, our data indicate that reference control spectra can vary on a week-to-week bassis despite adequate control of these variables. These data collectively suggest that the appropriateness of reusing reference controls depends broadly on the investigator’s experimental goals, and more specifically on the specific antibody-fluorochrome conjugates in the panel (and their fluorescence intensities. For example, in considering the variability in the emission spectra of PE-Cy7-CCR6 (**Online Figure 9B**), the variations of the highest magnitude (specifically variations where the absolute value of ΔNormalized MFI ≥ 0.02) are present in YG10, followed by B14, followed by B12-B13. Because the YG10 and B14 detectors correspond to the peak emissions of PE-Fire810 and RB780, respectively, it is likely that re-use of the PE-Cy7-CCR6 reference control will create unmixing errors between PE-Cy7 and either one or both of these fluorochromes. Thus, re-use of previously recorded PE-Cy7 reference controls is likely to result in unmixing errors in panels using PE-Fire810 or RB780. However, such an approach may be appropriate in panels lacking these fluorochromes. Given the use of both PE-Fire810 and RB780 in the current panel, these observations encouraged us to re-record specific reference controls (including PE-Cy7) with each experiment, beyond the originally planned monthly “Control Load” experiments.

**Figure S9.**
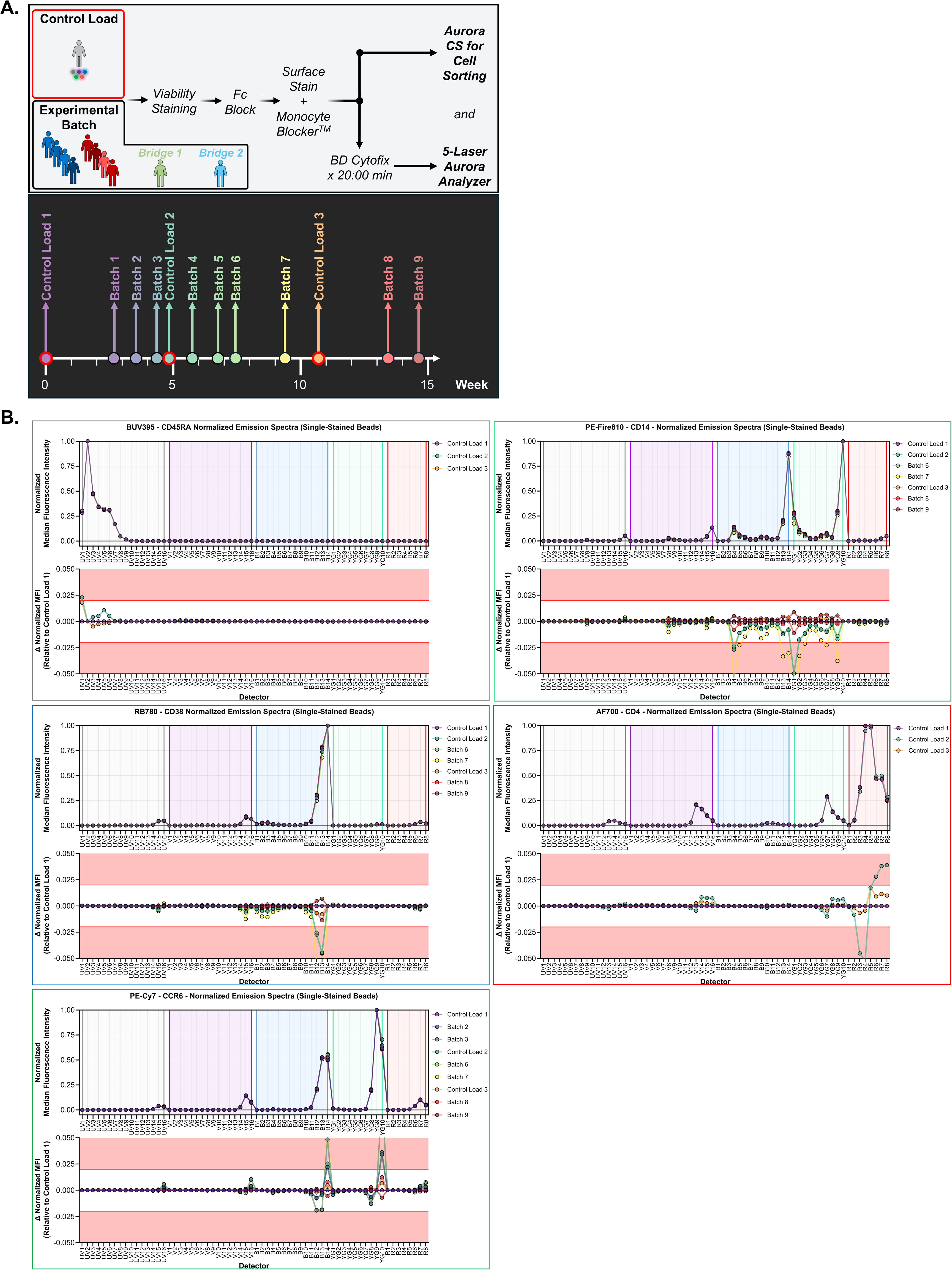
Application of this panel for simultaneous longitudinal immune profiling and cell sorting using the Cytek® Aurora Analyzer and Aurora CS. **A**. Schematic of the experimental approach for simultaneous longitudinal immune profiling and cell sorting using the Cytek^®^ Aurora Analyzer and Cytek^®^ Aurora CS, respectively. Experiments were performed over 15 weeks and are comprised of two “types” of experiments. “Control Load” experiments reflect the preparation of n=31 (including unstained cells) single-stained human PBMCs, n=29 (excluding the viability dye) single-stained Ultracomp eBeads^TM^ Compensation Beads, and n=1 fully stained PBMCs, for both the Cytek^®^ Aurora Analyzer and Cytek^®^ Aurora CS (total of n=61 samples per instrument). “Batch” experiments reflect the preparation of n=8-10 fully stained PBMCs from healthy donors (blue) and donors with a specific systemic autoimmune disease (red; data from these samples are not included in this manuscript). Each “Batch” experiment also includes two “Bridge” samples (light green, light blue), reflecting PBMC aliquots from two distinct healthy human donors which were included in all 9 “Batch” experiments. “Batch” experiments also included single-stained human PBMCs and single-stained beads for select antibody-fluorochrome conjugates. The staining procedure was executed identically for each experiment, with specific attention to (a) sample preparation; (b) temperature; (c) incubation times; (d) reagent lot number (including identical lot numbers for both tandem and non-tandem fluorochromes); (e) instrument selection, setup, and calibration. Samples were processed simultaneously on the 5-Laser Cytek^®^ Aurora Analyzer and Cytek^®^ Aurora CS. **B**. Normalized emission spectra for select antibody-fluorochrome conjugates (single-stained beads) recorded across multiple experimental timepoints, and quantification of the differences in the normalized emission spectra between the first recording of a given conjugate (“Control Load 1”) and subsequent recordings of the same conjugate. These data highlight the variability in the normalized emission spectra of reference controls that are prepared identically and analyzed based on identical acquisition settings, but recorded on different dates.

### UNBIASED ANALYSIS OF PANEL PERFORMANCE AND EXPERIMENTAL BATCH EFFECTS

In addition to manual inspection of the bivariate plots depicted in the gating strategy (**Figure 1** of the main text) and the NxN bivariate plot matrices of fully stained samples, we opted to leverage unbiased approaches to assess the panel’s performance at (1) a single point in time and (2) across multiple experimental batches (**Online Figure 10**). The analytical workflow was completed using the OMIQ cytometry analysis platform (https://omiq.ai) and is described in greater detail below. Briefly, this workflow was applied to unmixed .fcs files from a total of n=18 samples comprised of n=9 “Bridge 1” samples (Batches 1-9) and n=9 “Bridge 2” samples (Batches 1-9). Given the previous discussion regarding the pitfalls of re-use of previously recorded reference controls across multiple experimental batches, a small amount of spillover adjustment (generally 0-5%) was manually applied to each experimental batch. We then manually filtered the data to obtain live, CD45^+^ events. We subsequently generated UMAP projections of both normalized and non-normalized versions of these data. To compare the variation within a given sample (i.e., variation by experimental batch for a sample) to the variation between samples (i.e., variation by sample within a given experimental batch), we calculated pairwise Earth Mover’s Distances (EMDs) based on the UMAP coordinates for each sample.^4^ We quantified intra-sample variation by calculating the pairwise EMDs within each “Bridge Sample” across all 9 batches (ΔBridge1_Batches 2-9 vs. Batch 1_ and ΔBridge2_Batches 2-9 vs. Batch 1_).

We subsequently quantified inter-sample variation by calculating pairwise EMDs between Bridge 1 and Bridge 2 within a given experimental batch. These analyses indicate that the variation observed between distinct samples (i.e., Bridge 1 vs. Bridge 2; two healthy 20-25 year-old females) vastly exceeds the relatively minorl variation observed within a sample (Bridge_1_Batch_1 vs. Bridge_1_Batch_8), regardless of the computational normalization.

### DETAILED ANALYTICAL WORKFLOW

1. **<u>Scaling</u>**: Scaling was adjusted to ensure that the negative population for each parameter exhibited a unimodal distribution centered on a fluorescence intensity of 0.
2. **<u>Compensation</u>**: Because the normalized emission spectra of single-color controls exhibited slight, albeit nontrivial variations across experimental batches (**Online Figure 8**), a small amount of spillover matrix adjustment (0-5% for fluorochrome pairs with overlapping emission spectra) was applied to each experimental batch based on manual inspection of each batch’s NxN bivariate plot matrix (n=2 concatenated samples analyzed for each of the 9 experimental batches).
3. **<u>Gating</u>**: Data were then manually gated as depicted in **Figure 1** of the main text in order to identify single, live, CD45^+^ events (∼500,000 events/sample).
4. **<u>Subsampling the Cell Filter</u>**: Data were then subsampled to only these single, live, CD45^+^ events. The total number of events per sample was not modified.
5. **<u>[Optional] Normalization Across Experimental Batches</u>:** We used two different approaches to perform batch normalization (cyCombine^5^, CytoNorm^6^). These approaches are described in greater detail below, within this Step (5a). Nevertheless, we also sought to evaluate the longitudinal performance of the panel without any computational batch normalization. The analytical workflow for non-normalized data is also described below, within this Step (5b).

a. Normalization

i. FlowSOM: FlowSOM clustering was performed on all samples (n=18) using 29 of the 30 parameters in this panel (all except PerCP-CD45). Specific parameters include the following:

1. xdim: 12, ydim: 12, number of training iterations: 10, distance metric: euclidean, consensus metaclustering: k=30, random seed: 826
ii. cyCombine or CytoNorm: Two different normalization methods were independently applied to the data. The specific parameters for each normalization approach are listed here.

1. *cyCombine*: cyCombine was performed on all samples (n=18) using 29 of the 30 parameters in this panel (all except PerCP-CD45). Specific parameters include the following:

a. Batch Metadata: Experimental Batch, Anchor (optional): Bridge Sample 1; FlowSOM Metacluster k=30 used to group similar cells; parametric version of ComBat algorithm used for cyCombine normalization
2. *CytoNorm*: CytoNorm was performed on all samples (n=18) using 29 of the 30 parameters in this panel (all except PerCP-CD45). Specific parameters include the following:

a. Number of quantiles: 101, Batch Metadata Field: Experimental Batch, Control Metadata Field: Bridge Sample 1, Cluster Membership Feature: FlowSOM Metacluster k=30.
iii. Proceed to Step 6.
b. No normalization

No additional tasks are performed within this step. Proceed to Step 6.
6. **<u>Subsampling the Total Event Number Per Sample</u>**: Data from each sample (n=18 total) was downsampled to 100,000 live, CD45^+^ events per sample.
7. **<u>Dimensionality Reduction via UMAP</u>:** All files (n=18) were used to construct the UMAP projections. UMAP projections of normalized data (whether via cyCombine or CytoNorm) were constructed using all normalized parameters (29 total). UMAP projections of nonnormalized data were constructed using 29 of the 30 unmodified parameters (all parameters except PerCP-CD45). Additional settings/parameters to construct the UMAP were kept constant regardless of whether normalization had been implemented and are listed below:

a. Neighbors: 15, Minimum Distance: 0.4, Components: 2, Metric: euclidean, Learning Rate: 1, Epochs: 200, Random Seed: 6834, Embedding initialization: spectral
8. **<u>Visualization</u>**: UMAP projections and feature plots were subsequently generated.
9. **<u>Calculation of Earth Mover’s Distance (EMD)</u>**: The resulting UMAP coordinates for each event for each sample (total of 1.8 million values) were exported. Pairwise EMD values were calculated using R, as described previously.^4^

**Figure S10.**
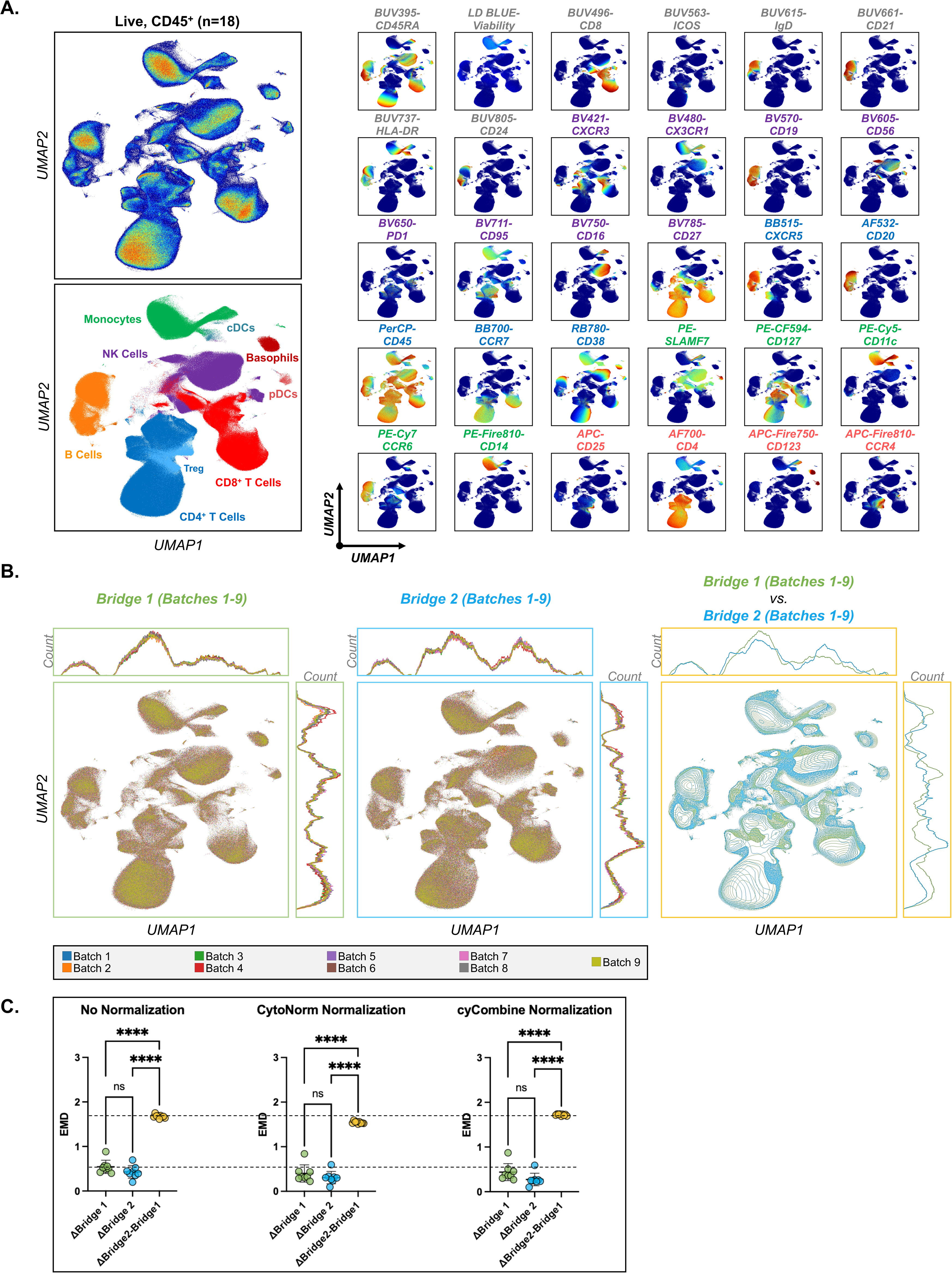
Longitudinal immune profiling reveals minimal batch effects even without computational normalization. **A.** UMAP projection of live, CD45^+^ events from all Bridge samples across all 9 experimental batches, generated using all parameters (n=29; normalized using cyCombine) except CD45- PerCP. Data from Bridge 1 (n=9) and Bridge 2 (n=9) samples across all batches were downsampled to include 100,000 live, CD45^+^ events per sample for a total of 1.8 million displayed events. Populations identified using the gating strategy depicted in Figure 1 of the main text were overlaid onto the UMAP. The expression profile of each parameter is depicted for all 30 parameters. **B.** Overlay of UMAP scatterplots and UMAP coordinate histograms by experimental batch for a given Bridge sample. **C.** Overlay of UMAP contour plots and UMAP coordinate histograms for Bridge Sample 1 (Batches 1-9, light green) and Bridge Sample 2 (Batches 1-9, light blue). **D.** Earth Mover’s Distance (EMD) of the UMAP coordinate distributions with or without computational normalization. “ΔBridge 1” reflects the EMD between Bridge Sample 1_Batch 1_ and Bridge Sample 1_Batches 2-9_. “ΔBridge 2” reflects the EMD between Bridge Sample 2_Batch 1_ and Bridge Sample 2_Batches 2-9_. “ΔBridge2-ΔBridge1” reflects the EMD between Bridge Sample 2 and Bridge Sample 1 of the same experimental batch. *,**,***,**** = *p* < 0.05, 0.01, 0.001, and 0.0001, respectively, via Brown-Forsythe and Welch ANOVA with adjustment for multiple comparisons using Dunnett’s T3 multiple comparisons test (assuming nonequal standard deviations). Comparisons which are not statistically significantly different are not depicted.

### APPLICATION OF THE PANEL TO THE AURORA CELL SORTER

All cell sorting experiments described here were performed on a standard 5-Laser Cytek^®^ Aurora CS using a 100-um nozzle. Daily QC and stream settings had been optimized by the staff of the Flow Cytometry Core at the Children’s Hospital of Philadelphia. No manual adjustments were made to the manufacturer-recommended assay settings (CytekAssaySetting) for each detector. Accuracy of aiming for 4-way sorting (Purity Mode) was verified manually. The drop-drive frequency and amplitude were adjusted to ensure a drop center of ∼260 and a drop interval of 20. Sort Monitoring was enabled. Autodrop Delay was then performed. Fully-stained samples were loaded on the instrument for 4-way or 6-way cell sorting via “Purity” mode. The event rate was maintained at ≤ 1/5 of the drop drive frequency (∼5000 events/second). Sort purity was verified by reloading a 10 uL aliquot of a sorted population to enable acquisition of ≤ 2000 total events.

The experimental approach (**Online Figure 9A**) enabled all experimental samples to be recorded on both the 5 Laser Aurora Analyzer and the Aurora CS. We therefore sought to evaluate the panel’s performance on the Aurora CS. Unbiased depiction of the data obtained on the Aurora CS appeared grossly similar to that of the Aurora Analyzer (**Online Figure 11**). We subsequently analyzed these data according to the gating strategy presented in **Figure 1**, and also included a direct comparison to the corresponding samples that were analyzed on the Aurora Analyzer (**Online Figure 12A**). Altogether, these data indicate excellent panel performance on the Aurora CS, with sufficient resolution to identify the immune subsets of interest. Our data also indicate that our approach can be used to sort immune cell subsets with high purity (≥90%) (**Online Figure 12B**). Notably, these purity estimates are likely an underestimate, as we sampled relatively few events (<2000 total) from the sorted populations.^7^ Nevertheless, these data verify the cross-platform capability of this panel. Sorted T cell subsets were activated *in vitro* using anti-CD3/CD28 Dynabeads^TM^ and processed for RNA fluorescence *in situ* hybridization and RNA sequencing.

**Figure S11.**
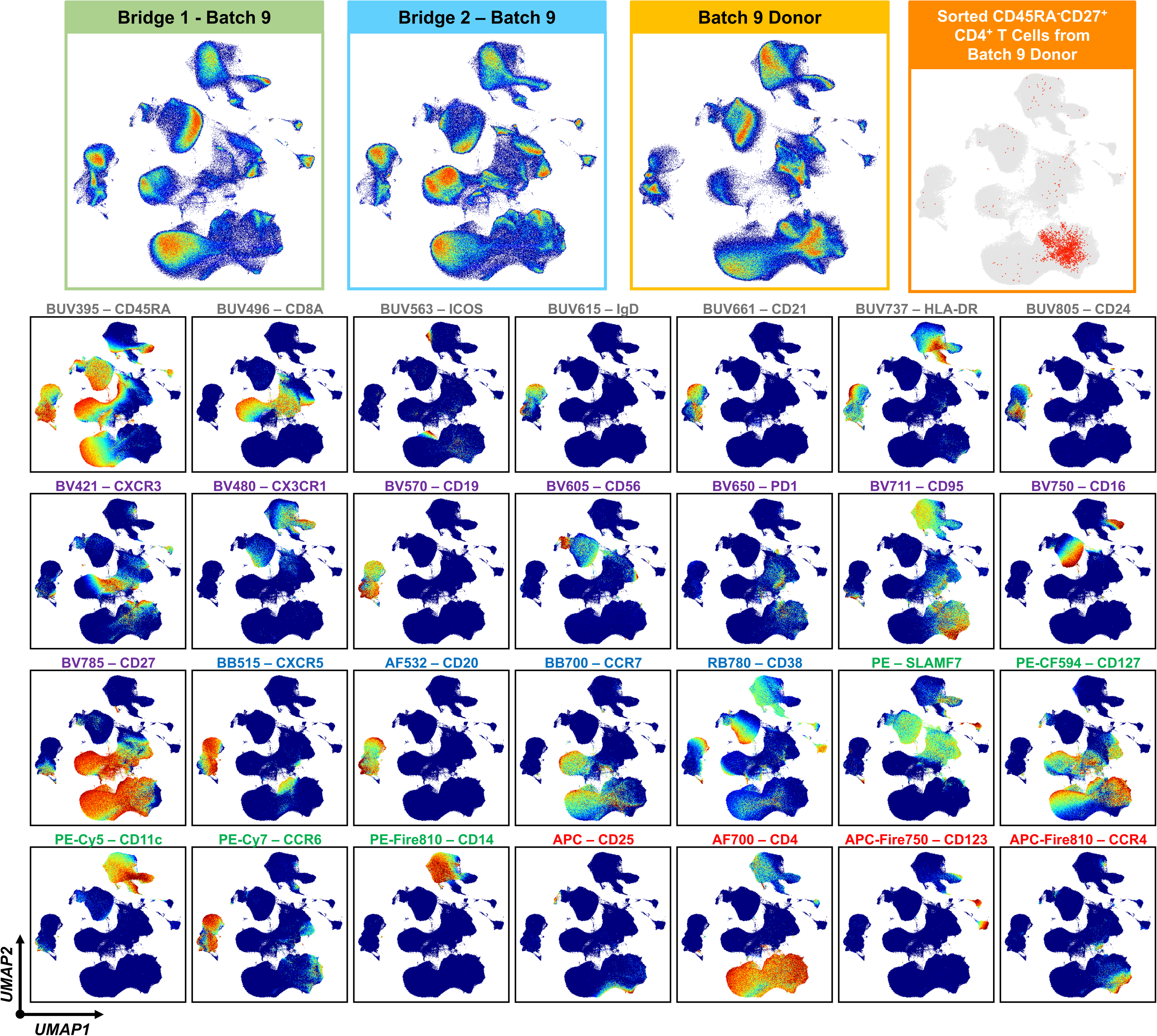
Dimensionality reduction of fully-stained samples analyzed using the Cytek^®^ Aurora CS. UMAP projections of live, CD45^+^ events from Bridge Sample 1_Batch 9_, Bridge Sample 2_Batch 9_, a unique sample from experimental Batch 9 (“Batch 9 Donor”), and sorted CD45RA^-^CD27^+^ CD4^+^ T cells from “Batch 9 Donor”, generated using all parameters (n=29) except CD45-PerCP. Data were downsampled to include 500,000 live, CD45^+^ events per sample; only 1,764 events were available for the sorted population from “Batch 9 Donor”. Colored-continuous scatterplots depict the marker expression profile. Data were acquired on the Cytek^®^ Aurora CS.

**Figure S12.**
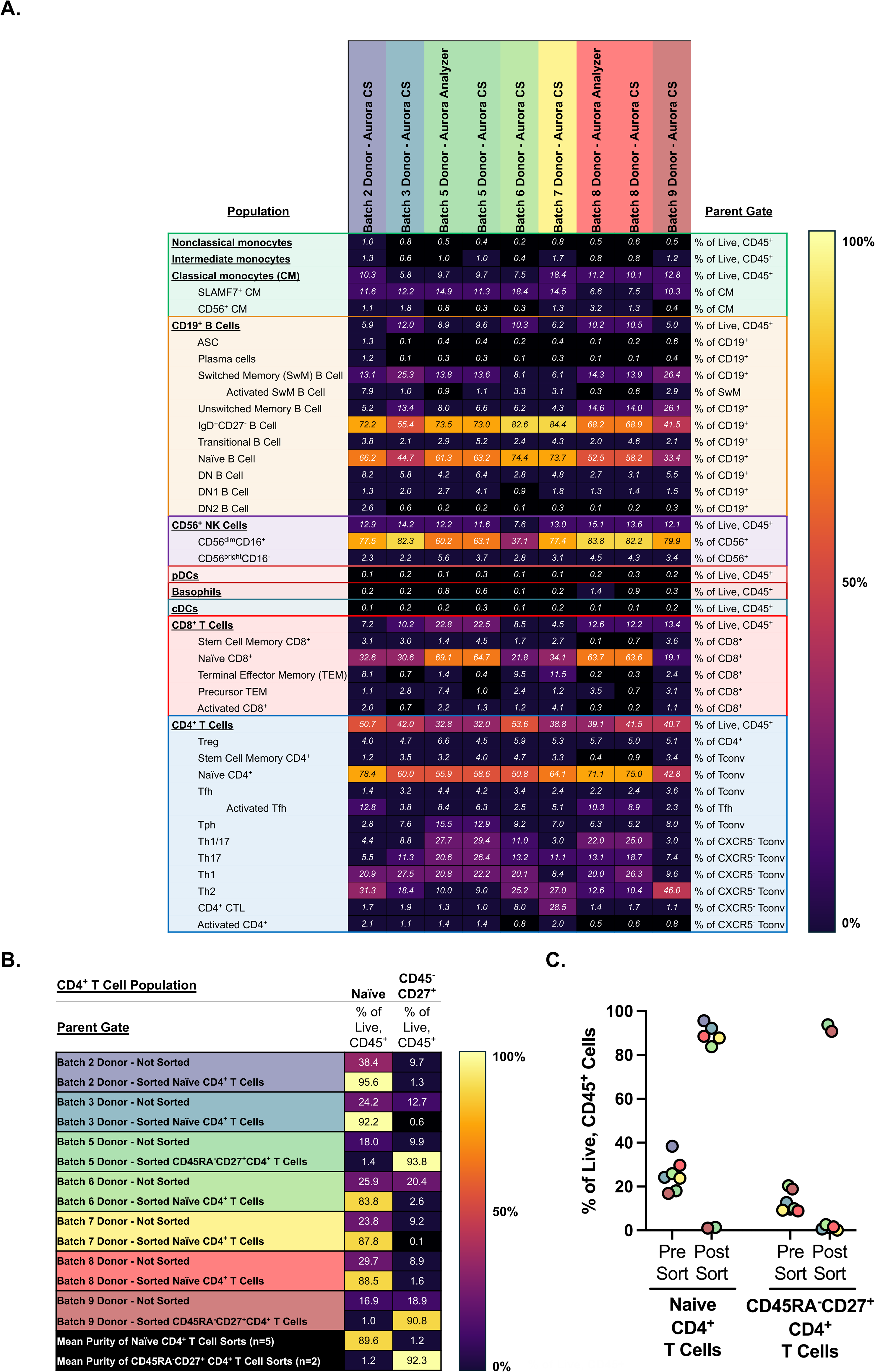
Quantification of immune cell subsets and post-sort purity on the Aurora CS. **A.** Quantification of immune cell subsets from distinct donors across multiple experimental batches using the gating strategy depicted in Figure 1 as determined via the Aurora CS. For two samples (Batch 5 Donor, Batch 8 Donor), the corresponding immune cell subsets quantified using the 5-Laser Aurora Analyzer are also provided. **B.** Assessment of post-sort purity for the 7 samples depicted in Panel A. Although four cell subsets were simultaneously purified from each donor, the purity assessment was performed using only the most abundant sorted subset. Purified naïve CD4^+^ T cells were used to assess the post-sort purity for 5 of the 7 samples. Purified central/early effector memory (CD45RA^-^CD27^+^) CD4^+^ T cells were used to assess the post-sort purity for the remaining 2 of 7 samples. To preserve sorted cells for downstream applications, <2,000 events were collected for each purity assessment. Data are presented as a heatmap and **C.** in graphical format. Circles are colored according to the experimental batch.

### CROSS REFERENCES TO RELATED PANELS

Cross references to related panels are provided in **Table 1** and elaborated on in the Main Text.

### STAINING PROTOCOL

The following detailed protocol can be used to prepare fully stained PBMC samples and single-color reference controls (cells and beads) for analysis on the 5 Laser Cytek® Aurora Analyzer and cell sorting on the Aurora CS.

#### Reagents/Materials

- Peripheral Blood Mononuclear Cell (PBMC) Media*

- 10% heat-inactivated fetal bovine serum in RPMI
- 1% sodium pyruvate
- 1% GlutaMax
- 1% Pen/Strep
- Storage: 4°C
- Shelf-Life: Within 1 month of preparation
- Calcium/magnesium-free phosphate-buffered saline (PBS)

- Storage: Room temperature
- Shelf-Life: Expiration date
- Trypan Blue Solution, 0.4% (ThermoFisher Scientific Catalog #: 15250061)

- Storage: Room temperature
- Shelf-Life: Expiration date
- FACS Buffer*

- 0.5% bovine serum albumin in calcium/magnesium-free phosphate-buffered saline (if sorting, filter through 0.22 μm filter)
- Storage: 4°C
- Shelf-Life: Within 1 month of preparation
- Live/Dead Fixable BLUE

- Lyophilized contents were resuspended in DMSO according to the manufacturer’s instructions
- Storage: −20°C
- Shelf-Life: Expiration date
- Antibody-fluorochrome conjugates

- Storage: 4°C, dark
- Shelf-Life: Expiration date
- BD Horizon Brilliant Stain Buffer Plus (BD Biosciences, Catalog #: 566385)

- Storage: 4°C, dark
- Shelf-Life: Expiration date
- Human TruStain FcX (BioLegend, Catalog #: 426101-3)

- Storage: 4°C, dark
- Shelf-Life: Expiration date
- TrueStain Monocyte Blocker (BioLegend, Catalog #: 426101-3)

- Storage: 4°C, dark
- Shelf-Life: Expiration date
- [Optional] Ultracomp eBeads 2222-41)

- Storage: 4°C, dark
- Compensation Beads (Invitrogen, Catalog number: 01-
- Shelf-Life: Expiration date
- SpectroFlo® QC Beads (Cytek, Catalog number: SKU B7-10001)

- Storage: 4°C, dark
- Shelf-Life: Expiration date

#### Staining Protocol

All staining is performed in 96-well polypropylene round-bottom plates.

1. **Antibody Master Mix (AMM) “base” preparation**: *Given the number of fluorochromes included in this panel, the AMM “base” is prepared in advance of cell thawing to ensure timely addition of the complete antibody master mix to thawed cells during the staining process. Estimated time: ∼60 minutes.*

1.1 *<u>Antibody mixing and aggregate removal</u>*

1.1.1 All antibody-fluorochrome conjugates are briefly vortexed for several seconds (**Online Table 2**). Brilliant ultraviolet (BUV), brilliant violet (BV), and brilliant blue (BB) fluorochromes are then centrifuged at 16,000 g x 5 min at 4°C in order to minimize antibody/fluorochrome aggregates.^3^ The antibody-fluorochrome tubes are then placed on ice until further use.
1.2 *<u>Preparation of “predilutions” of dilute antibody-fluorochrome pairs</u>*

1.2.1 Several antibody-fluorochrome pairs in this panel are used at concentrations < 1:800. If staining only 1-3 samples, the total required volume of AMM is quite small, thus requiring the transfer of very small volumes of these “dilute” antibody-fluorochrome conjugates. To circumvent the potential imprecision associated with pipetting small volumes across multiple experiments, we prepare “pre-dilutions” for those antibody-fluorochrome conjugates that are particularly dilute. This approach ensures that the minimum volume of antibody-fluorochrome conjugate is 2 uL. These calculations are performed before the start of the experiment as depicted below (**Online Table 3**). Note that this approach is not necessary as the number of samples being stained increases.
1.3 *<u>Preparation of base antibody surface stain master mix:</u>*

1.3.1 Transfer x uL of Brilliant Stain Buffer PLUS, where x represents 25% of the total desired volume of Master Mix that will be prepared (50 uL of Master Mix is used for every 5 million cells stained), to a 1.5 mL Eppendorf tube labeled “AMM Base.”
1.3.2 Transfer the required volume of each antibody-fluorochrome conjugate.
1.3.3 Store the “AMM Base” tube at 4°C, in the dark
2. **Preparation of working concentrations for single-stained controls:** *Single-stained cells are used as reference controls for the majority of antibody-fluorochrome pairs in this panel. Estimated time: ∼30 minutes.*

2.1 Obtain 1.5 mL Eppendorf tubes (n=29) and label them according to the identity of the fluorochromes in the panel.
2.2 If cellular reference controls are desired (most appropriate for the results described here), prepare 75 uL of the specified dilution for each antibody-fluorochrome pair.
2.3 If both cellular and bead-based reference controls are desired, prepare 150 uL of the specified dilution for each antibody-fluorochrome pair.
2.4 Once all control solutions have been prepared, store them at 4°C in the dark.
3. **Cryopreserved PBMC Sample Thawing and Preparation for Staining:** *The thawing process is not trivial and has tremendous potential to influence post-thaw yield and viability. Estimated time: ∼10 minutes for 2 samples.*

3.1 *<u>Sample Thawing</u>*

3.1.1 Suspend up to two PBMC cryovials in a 37°C waterbath for 2:00 minutes exactly (assuming 1 mL of volume per tube; duration may differ based on volume per cryovial). At the 2:00 mark, suspend the tubes for up to an additional 30 seconds, but taking care to remove the tubes when only a small ice crystal remains. At this time, transfer the contents of the cryovial to a sterile 15 mL conical tube. Obtain pre-warmed PBMC media and, using a 10 mL serological pipette, draw up 11 mL of PBMC media and slowly dispense 1 mL (dropwise; about 1 mL every 3 seconds) into the now empty cryovial (wash), and immediately begin dispensing 5 mL into the 15 mL conical tube containing the cell suspension. Once 5 mL of media have been transferred, the remaining 5 mL can be added more quickly (∼1 mL every 2 seconds).
3.1.2 Once all of the media has been added, draw up 3-5 mL of the now diluted cell suspension, and dispense this volume gently to mix. The DMSO within the PBMC cryopreservation medium will now disperse throughout the total 10 mL volume. Failure to perform this gentle mixing could result in a 10-15% reduction in cell viability.
3.1.3 Centrifuge the cell suspension at 500 *g* for 5 minutes.
3.2 *<u>Cell Plating</u>*

3.2.1 Following the centrifugation, resuspend the cell pellet in x mL of warm media, where x is one fifth of the anticipated cell count, in millions of cells. Count the cell suspension by performing a 1:5 dilution of a sample aliquot using PBS, and then a 1:1 dilution using Trypan Blue (total 10X dilution). Allow the stock cell suspension to rest on ice.
3.2.2 After counting each sample, bring each conical tube up to 10 mL of total volume using PBS, and centrifuge the samples at 400 g for 5 minutes.
3.2.3 Following the centrifugation, resuspend each sample in a sufficient volume to obtain a total cell concentration of 25 million cells/mL. Transfer 200 uL of each sample to the wells of a round-bottom 96-well plate. If necessary, pool cells from multiple samples at this time and transfer anywhere from 50-200 uL of the pooled sample into a total of 31 wells of the 96-well plate. These wells will be used as reference controls (single-stained cells), including reference controls for unstained cells.
3.2.4 Centrifuge the 96-well plate at 400 g for 5 minutes and thaw an aliquot of Live-Dead Fixable Blue during this time.
4. **Staining:** *Estimated time: ∼70-90 minutes; all steps performed at room temperature*

4.1 *<u>Viability Dye</u>*

4.1.1 During the centrifugation of the cell plate, thaw an aliquot of Live-Dead Fixable Blue and prepare a 1:400 dilution in PBS (room temperature). Ensure the preparation of a sufficient volume to stain all samples and the single viability dye reference control.
4.1.2 Following the centrifugation of the cell plate, quickly gently flick out the supernatant, taking care to ensure that there is no residual volume in the cell wells.
4.1.3 Transfer 50 uL of the 1:400 Live-Dead Fixable Blue solution to each sample well and to the corresponding reference control well. Add 50 uL of room-temperature PBS to the remaining single-color reference control wells.
4.1.4 Cover the plate with aluminum foil and incubate it for 15:00 minutes at room temperature.
4.1.5 At this time, remove both the AMM base and the single-color reference control solutions from 4°C, and place them at room temperature in the dark so that they can begin to equilibrate to room temperature.
4.1.6 Following the viability dye incubation, transfer 175 uL of cold FACS buffer to all wells (total 225 uL per well).
4.1.7 Centrifuge the 96-well plate at 400 *g* for 5 minutes.
4.1.8 NOTE: FACS Buffer should now be stored at room temperature for all subsequent steps.
4.2 *<u>Fc Block</u>*

4.2.1 During this centrifugation step, prepare a 1:100 solution of TruStain Fc block in a sufficient volume of FACS Buffer such that 50 uL of this solution can be added to every cell-containing well.
4.2.2 Following the centrifugation, quickly gently flick out the supernatant, taking care to ensure that there is no residual volume in the cell wells. Quickly transfer 50 uL of the Fc block solution (1:100) to all cell-containing wells.

4.2.2.1 NOTE: Use of an additional 96-well plate to contain sufficient volumes of the solution being added may facilitate the rapid addition of these solutions via multichannel pipetting.
4.2.3 Cover the plate with aluminum foil and incubate it for 15:00 minutes at room temperature.

4.2.3.1 At this time, transfer the remaining components of the AMM base in order to generate the full Antibody Master Mix. This requires the addition of the specified volume of TruStain Monocyte Blocker (final concentration 1:75) and FACS buffer, which will generate the total desired volume of AMM.
4.2.4 Following the Fc block incubation, transfer 175 uL of FACS buffer to all wells (total 225 uL per well).
4.2.5 *<u>If bead-based reference controls are also desired:</u>*

4.2.5.1 Vortex a vial of compensation beads and transfer 0.5 drops of compensation beads to 29 wells (to generate 29 reference controls). This can be achieved by generating a suspension of n/2 drops (n = the number of reference control samples). Transfer 175 uL of FACS buffer to all bead-containing wells.
4.2.6 Centrifuge the 96-well plate at 400 *g* for 5 minutes.
4.3 *<u>Antibody Master Mix (Surface Staining Only)</u>*

4.3.1 During this centrifugation step, obtain a new, sterile 96-well round bottom plate (“Stain Plate”) and transfer 65 uL of gently mixed, now complete AMM to the specific wells corresponding to the fully-stained samples on the cell plate. Similarly, transfer 65 uL of each single-stained reference control (or FACS buffer for “Unstained” and “Live/Dead” reference controls) to the specific wells corresponding to the single-color controls on the cell plate.
4.3.2 Following the centrifugation, quickly gently flick out the supernatant, taking care to ensure that there is no residual volume in the cell wells. Using a multichannel pipette, quickly transfer 50 uL of solution from the corresponding wells of the “Stain Plate” to the cell plate and gently mix all samples.
4.3.3 Cover the cell plate with aluminum foil and incubate it for 40:00 minutes at room temperature.
4.3.4 At this time, place the FACS buffer back on ice.
4.3.5 Following the AMM incubation, transfer 175 uL of now cool FACS buffer to all wells (total 225 uL per well).
4.3.6 <u>PAUSE</u>:

4.3.6.1 If analysis on the 5-laser Cytek^®^ Aurora Analyzer *<u>or</u>* cell sorting on the Cytek^®^ Aurora CS is the sole experimental goal (i.e., either one or the other, but not both), proceed to the next centrifugation step.
4.3.6.2 If the investigator wishes to prepare samples simultaneously for *<u>both</u>* cell sorting on the Cytek^®^ Aurora CS and analysis on the 5-laser Cytek^®^ Aurora Analyzer, then proceed to the following step:

4.3.6.2.1 The goal of this step is to transfer representative fully stained samples and reference controls to a new plate that will be used for analysis on the 5-laser Cytek^®^ Aurora Analyzer. Assuming that the majority of a fully stained sample will be processed for cell sorting, transfer 60-100 μL of each fully stained unique sample to the corresponding wells of a new 96-well U-bottom plate labeled “Analyzer”. Transfer 100 uL of all reference controls to the corresponding wells of this new plate as well. Top off all wells across both plates with FACS buffer to achieve a total volume of 200 uL.
4.3.7 Centrifuge the 96-well plate(s) at 400 *g* for 5 minutes.
5. **Fixation for Analysis or Resuspension in FACS for Cell Sorting or Both:** *Estimated time: 5 minutes (cell sorting); 30 minutes (cell analysis)*

5.1 Following the centrifugation, quickly and gently flick out the supernatant, taking care to ensure that there is no residual volume in the cell wells.

5.1.1 <u>If cells are being prepared for cell sorting</u>, transfer 200 uL of cool FACS buffer to all wells containing cells or beads.

5.1.1.1 Cover the cell plate and store on ice in anticipation of cell sorting.
5.1.2 <u>If cells are being prepared for analysis</u>, transfer 200 uL of cold (4°C) BD Cytofix buffer to all wells containing either cells or beads.

5.1.2.1 Cover the cell plate with aluminum foil and incubate it for 20:00 minutes at room temperature.
5.1.2.2 Following the fixative incubation, centrifuge the 96-well plate at 400 *g* for 5 minutes.
5.1.2.3 Following the centrifugation, quickly and gently flick out the supernatant, taking care to ensure that there is no residual volume in the wells.
5.1.2.4 Transfer 200 uL of cool FACS buffer to all wells containing either cells or beads.
5.1.2.5 Cover the cell plate and store on ice in anticipation of analysis (in this manuscript, samples were recorded within 4 hours of completion of staining; it may be permissible to analyze the samples after this time, though the marker resolution may differ subtly from the results reported here).

---

**Online Table S2.**
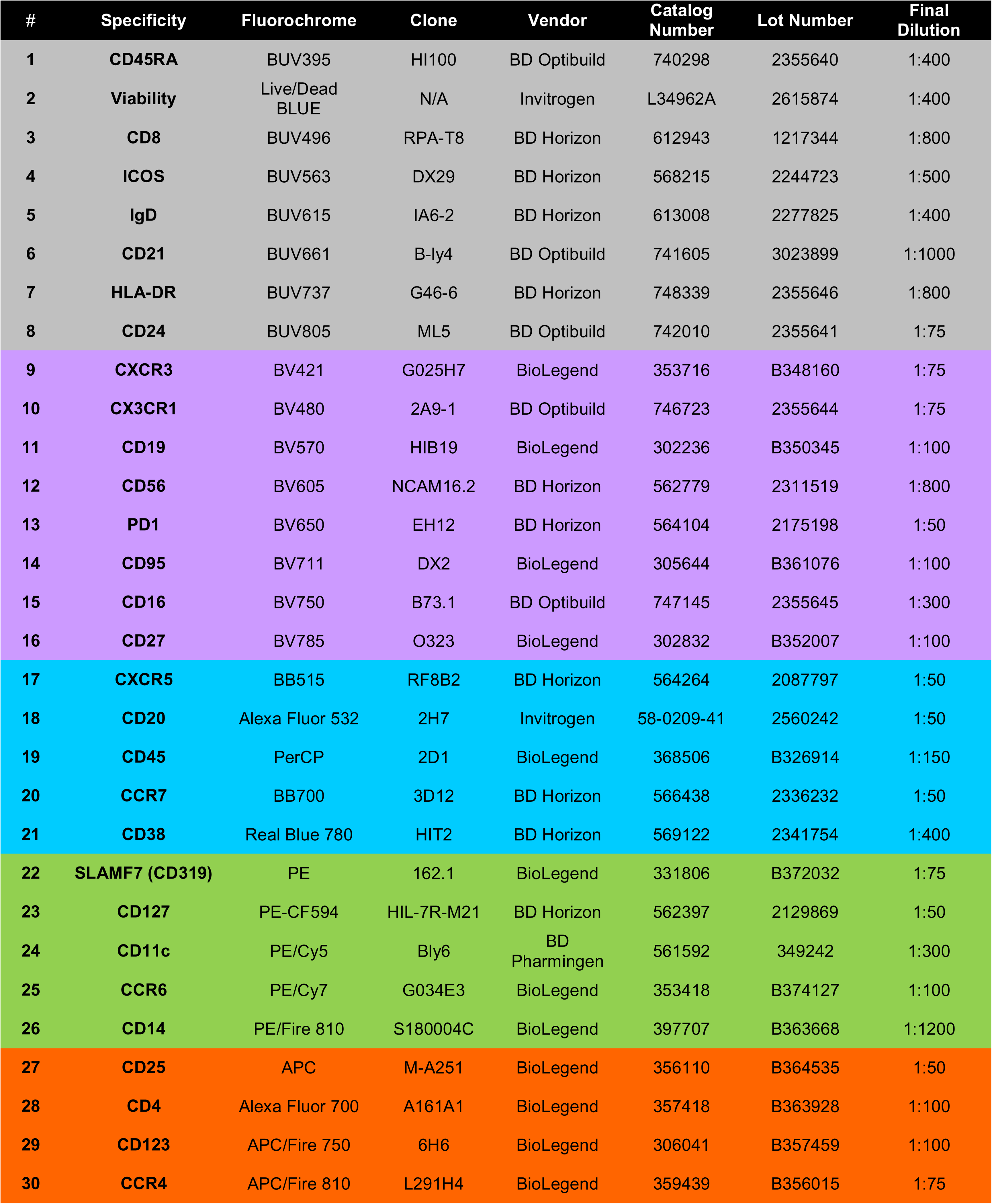
Reagent Information.

**Online Table 2. Reagent Information**
| # | Specificity | Fluorochrome | Clone | Vendor | Catalog Number | Lot Number | Final Dilution |
| --- | --- | --- | --- | --- | --- | --- | --- |
| 1 | CD45RA | BUV395 | HI100 | BD Optibuild | 740298 | 2355640 | 1:400 |
| 2 | Viability | Live/Dead BLUE | N/A | Invitrogen | L34962A | 2615874 | 1:400 |
| 3 | CD8 | BUV496 | RPA-T8 | BD Horizon | 612943 | 1217344 | 1:800 |
| 4 | ICOS | BUV563 | DX29 | BD Horizon | 568215 | 2244723 | 1:500 |
| 5 | IgD | BUV615 | IA6-2 | BD Horizon | 613008 | 2277825 | 1:400 |
| 6 | CD21 | BUV661 | B-ly4 | BD Optibuild | 741605 | 3023899 | 1:1000 |
| 7 | HLA-DR | BUV737 | G46-6 | BD Horizon | 748339 | 2355646 | 1:800 |
| 8 | CD24 | BUV805 | ML5 | BD Optibuild | 742010 | 2355641 | 1:75 |
| 9 | CXCR3 | BV421 | G025H7 | BioLegend | 353716 | B348160 | 1:75 |
| 10 | CX3CR1 | BV480 | 2A9-1 | BD Optibuild | 746723 | 2355644 | 1:75 |
| 11 | CD19 | BV570 | HIB19 | BioLegend | 302236 | B350345 | 1:100 |
| 12 | CD56 | BV605 | NCAM16.2 | BD Horizon | 562779 | 2311519 | 1:800 |
| 13 | PD1 | BV650 | EH12 | BD Horizon | 564104 | 2175198 | 1:50 |
| 14 | CD95 | BV711 | DX2 | BioLegend | 305644 | B361076 | 1:100 |
| 15 | CD16 | BV750 | B73.1 | BD Optibuild | 747145 | 2355645 | 1:300 |
| 16 | CD27 | BV785 | O323 | BioLegend | 302832 | B352007 | 1:100 |
| 17 | CXCR5 | BB515 | RF8B2 | BD Horizon | 564264 | 2087797 | 1:50 |
| 18 | CD20 | Alexa Fluor 532 | 2H7 | Invitrogen | 58-0209-41 | 2560242 | 1:50 |
| 19 | CD45 | PerCP | 2D1 | BioLegend | 368506 | B326914 | 1:150 |
| 20 | CCR7 | BB700 | 3D12 | BD Horizon | 566438 | 2336232 | 1:50 |
| 21 | CD38 | Real Blue 780 | HIT2 | BD Horizon | 569122 | 2341754 | 1:400 |
| 22 | SLAMF7 (CD319) | PE | 162.1 | BioLegend | 331806 | B372032 | 1:75 |
| 23 | CD127 | PE-CF594 | HIL-7R-M21 | BD Horizon | 562397 | 2129869 | 1:50 |
| 24 | CD11c | PE/Cy5 | Bly6 | BD Pharmingen | 561592 | 349242 | 1:300 |
| 25 | CCR6 | PE/Cy7 | G034E3 | BioLegend | 353418 | B374127 | 1:100 |
| 26 | CD14 | PE/Fire 810 | S180004C | BioLegend | 397707 | B363668 | 1:1200 |
| 27 | CD25 | APC | M-A251 | BioLegend | 356110 | B364535 | 1:50 |
| 28 | CD4 | Alexa Fluor 700 | A161A1 | BioLegend | 357418 | B363928 | 1:100 |
| 29 | CD123 | APC/Fire 750 | 6H6 | BioLegend | 306041 | B357459 | 1:100 |
| 30 | CCR4 | APC/Fire 810 | L291H4 | BioLegend | 359439 | B356015 | 1:75 |

**Online Table S3.**
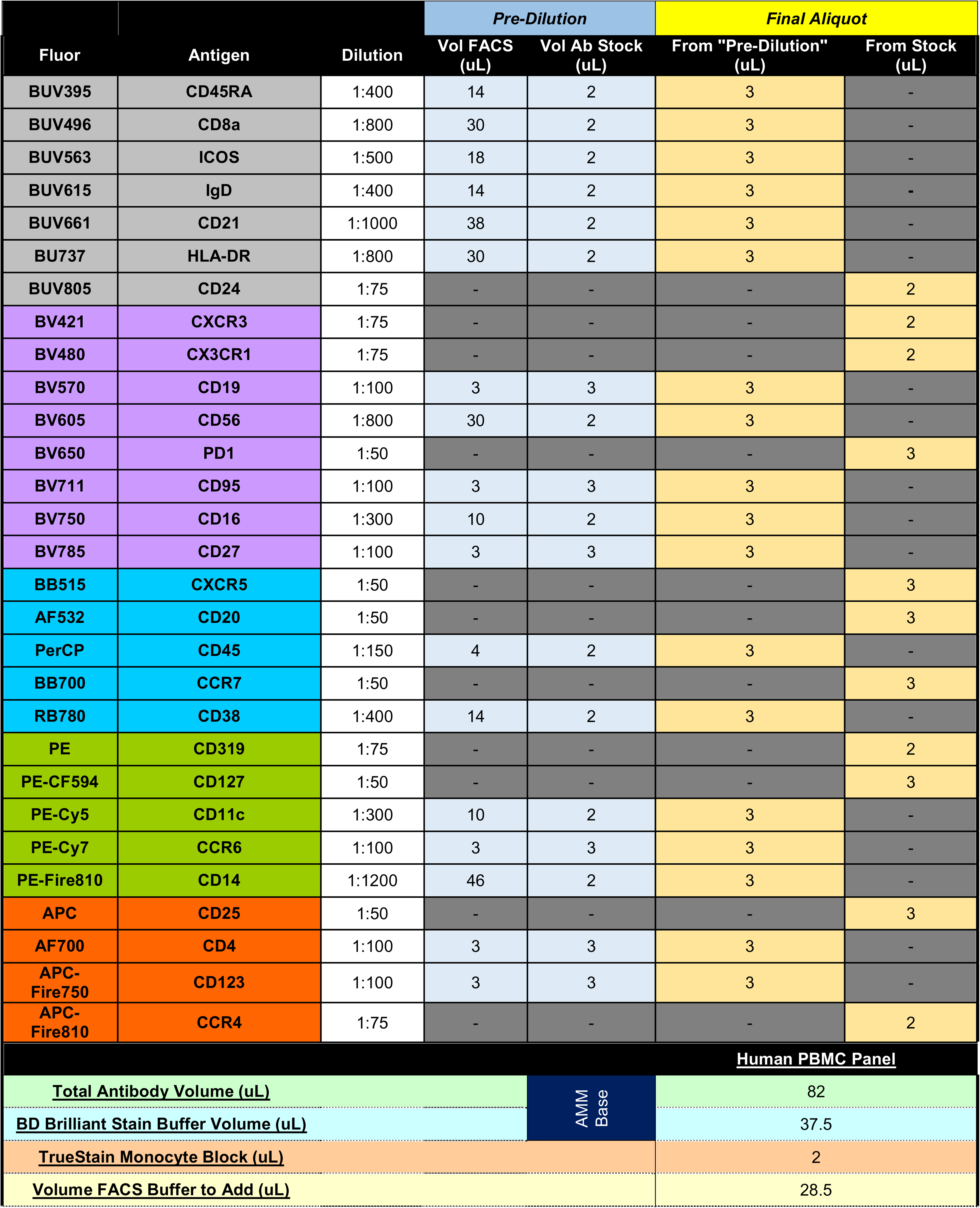
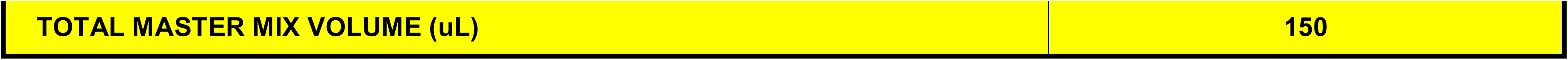
Example of the approach used to prepare small volumes of antibody master mix. Pre-dilutions for dilute conjugates are initially prepared as depicted above. The desired antibody master mix is then prepared using either the prepared “pre-dilutions” or the antibody stock.

|  |  |  | Pre-Dilution |  | Final Aliquot |  |
| --- | --- | --- | --- | --- | --- | --- |
| Fluor | Antigen | Dilution | Vol FACS (uL) | Vol Ab Stock (uL) | From "Pre-Dilution" (uL) | From Stock (uL) |
| BUV395 | CD45RA | 1:400 | 14 | 2 | 3 | - |
| BUV496 | CD8a | 1:800 | 30 | 2 | 3 | - |
| BUV563 | ICOS | 1:500 | 18 | 2 | 3 | - |
| BUV615 | IgD | 1:400 | 14 | 2 | 3 | - |
| BUV661 | CD21 | 1:1000 | 38 | 2 | 3 | - |
| BU737 | HLA-DR | 1:800 | 30 | 2 | 3 | - |
| BUV805 | CD24 | 1:75 | - | - | - | 2 |
| BV421 | CXCR3 | 1:75 | - | - | - | 2 |
| BV480 | CX3CR1 | 1:75 | - | - | - | 2 |
| BV570 | CD19 | 1:100 | 3 | 3 | 3 | - |
| BV605 | CD56 | 1:800 | 30 | 2 | 3 | - |
| BV650 | PD1 | 1:50 | - | - | - | 3 |
| BV711 | CD95 | 1:100 | 3 | 3 | 3 | - |
| BV750 | CD16 | 1:300 | 10 | 2 | 3 | - |
| BV785 | CD27 | 1:100 | 3 | 3 | 3 | - |
| BB515 | CXCR5 | 1:50 | - | - | - | 3 |
| AF532 | CD20 | 1:50 | - | - | - | 3 |
| PerCP | CD45 | 1:150 | 4 | 2 | 3 | - |
| BB700 | CCR7 | 1:50 | - | - | - | 3 |
| RB780 | CD38 | 1:400 | 14 | 2 | 3 | - |
| PE | CD319 | 1:75 | - | - | - | 2 |
| PE-CF594 | CD127 | 1:50 | - | - | - | 3 |
| PE-Cy5 | CD11c | 1:300 | 10 | 2 | 3 | - |
| PE-Cy7 | CCR6 | 1:100 | 3 | 3 | 3 | - |
| PE-Fire810 | CD14 | 1:1200 | 46 | 2 | 3 | - |
| APC | CD25 | 1:50 | - | - | - | 3 |
| AF700 | CD4 | 1:100 | 3 | 3 | 3 | - |
| APC-Fire750 | CD123 | 1:100 | 3 | 3 | 3 | - |
| APC-Fire810 | CCR4 | 1:75 | - | - | - | 2 |
| Human PBMC Panel |  |  |  |  |  |  |
| Total Antibody Volume (uL) |  |  |  | AMM Base | 82 |  |
| BD Brilliant Stain Buffer Volume (uL) |  |  |  |  | 37.5 |  |
| TrueStain Monocyte Block (uL) |  |  |  |  | 2 |  |
| Volume FACS Buffer to Add (uL) |  |  |  |  | 28.5 |  |

|  |  |
| --- | --- |
| TOTAL MASTER MIX VOLUME (uL) | 150 |

